# Genetic basis of enhanced stress resistance in long-lived mutants highlights key role of innate immunity in determining longevity

**DOI:** 10.1101/2021.07.19.452975

**Authors:** Sonja K. Soo, Paige D. Rudich, Meeta Mistry, Jeremy M. Van Raamsdonk

## Abstract

Mutations that extend lifespan are associated with enhanced resistance to stress. To better understand the molecular mechanisms underlying this relationship, we studied nine long-lived *C. elegans* mutants representative of different pathways of lifespan extension. We directly compared the magnitude of their lifespan extension and their ability to resist various external stressors (heat, oxidative stress, bacterial pathogens, osmotic stress, and anoxia). Furthermore, we analysed gene expression in each of these mutants to identify genes and pathways responsible for the enhanced resistance to stress. All of the examined long-lived mutants have increased resistance to one or more type of stress. Resistance to each of the examined types of stress had a significant, positive correlation with lifespan, with bacterial pathogen resistance showing the strongest relationship. All of the examined long-lived mutants have significant upregulation of multiple stress response pathways but differ in which stress response pathway has the greatest enrichment of genes. We used RNA sequencing data to identify which genes are most highly correlated with each type of stress resistance. There was a highly significant overlap between genes highly correlated with stress resistance, and genes highly correlated with longevity, suggesting that the same genetic pathways drive both phenotypes. This was especially true for genes correlated with bacterial pathogen resistance, which showed an 84% overlap with genes correlated with lifespan. Overall, our results demonstrate a strong correlation between stress resistance and longevity that results from the high degree of overlap in genes contributing to each phenotype.

**SIGNIFICANCE STATEMENT:** While increased resistance to stress has been correlated with longevity, the genetic basis for this relationship is incompletely understood. To advance our understanding of the relationship between stress resistance and lifespan, we measured lifespan, stress resistance and gene expression in a panel of nine long-lived mutants in *C. elegans*. All of the long-lived mutants exhibit enhanced resistance to at least one external stressor resulting from significant upregulation of multiple stress response pathways. Importantly, our data indicates that the same genetic pathways control stress resistance and lifespan, thereby accounting for the strong correlation between these two phenotypes. This work demonstrates the importance of innate immune signaling and other stress response pathways in determining longevity.

## INTRODUCTION

Paradigm-shifting work in the worm *C. elegans* has identified single gene mutations that significantly extend lifespan, thereby demonstrating a clear contribution of genetics to lifespan determination. Since the first genes to increase lifespan were identified in *C. elegans* (1–4), single gene mutations have also been shown to extend lifespan in other model organisms including yeast, flies and mice (5–7). Importantly, many of these lifespan-extending genes or interventions are evolutionarily conserved (8).

The availability of genetic mutants with extended lifespan has facilitated investigation into the mechanisms underlying their increased longevity, and the categorization of genetic mutants into specific pathways of lifespan extension. These pathways include: decreased insulin-IGF1 signaling (2, 4), mild impairment of mitochondrial function (9–11), dietary restriction (12), germ line inhibition (13), reduced chemosensation (14), decreased translation (15, 16), and increased mitochondrial reactive oxygen species (ROS) (17).

In exploring factors contributing to longevity, it has been observed that many long-lived mutants exhibit increased resistance to at least one type of external stressor. The best characterized example is the long-lived insulin-IGF1 receptor mutant *daf-2*, which has increased resistance to oxidative stress, heat stress, osmotic stress, anoxia, heavy metals, and bacterial pathogens (18–22). However, many counterexamples also exist. Mutations in the mitochondrial superoxide dismutase gene (*sod-2*) increase lifespan but decrease resistance to oxidative stress (17). The inverse also occurs where mutations that increase stress resistance result in decreased lifespan; such as the constitutive activation of the mitochondrial unfolded protein response (mitoUPR) transcription factor ATFS-1 (23–25). Decreasing stress resistance in *daf-2* mutants through disruption of *gpdh-1/2* or *nhl-1* diminishes stress resistance but increases lifespan (22). Thus, although stress resistance and lifespan are correlated, they can be experimentally dissociated.

A relationship between stress resistance and aging is also supported by the observation that resistance to multiple external stressors declines with age (26), at least in part due to a genetically-programmed downregulation of stress response pathways (27, 28). Importantly, a positive relationship between stress resistance and lifespan is conserved across species (29–31). Resistance to physiological stress is proposed to be one of the eight hallmarks of aging (32).

In this study, we measured stress resistance in nine long-lived mutants which represent multiple different pathways of lifespan extension and compared the stress resistance of each mutant to the mutation-induced lifespan extension and changes in gene expression across the genome. By quantifying all of these factors within the same study, we were able to directly compare lifespan and stress resistance across all of the long-lived mutants and examine the genetic underpinnings of stress resistance in these long-lived strains. We found that all nine of the long-lived mutants that we examined have increased resistance to at least one external stressor, and that all six of the examined types of stress resistance are significantly correlated with lifespan. In exploring the underlying mechanisms, we found that all of the long-lived mutants exhibit upregulation of genetic targets of multiple stress response pathways. Finally, we find that genes correlated with stress resistance exhibit a highly significant overlap with genes correlated with lifespan. Overall, this work advances our understanding of the relationship between stress resistance and longevity and demonstrates that the genetic pathways that contribute to these two phenotypes are highly overlapping.

## RESULTS

### Long-lived mutants show different magnitudes of lifespan extension

To determine the extent to which different types of stress resistance are correlated with longevity, and which types of stress resistance exhibit the highest correlation, we quantified resistance to stress and lifespan in nine long-lived mutants representing different pathways of lifespan extension. Measuring the stress resistance of these mutants together in the same assay allowed us to compare the relative magnitude of stress resistance with the lifespan extension of each mutant.

The mutants that we examined included: *daf-2* worms, which have decreased insulin-IGF1 signalling (4); *eat-2* worms, which are a model of dietary restriction (12); *ife-2* worms, which have decreased translation (15, 16); *clk-1, isp-1* and *nuo-6* worms, which have mild impairment of mitochondrial function (3, 9–11); *sod-2* worms, which have increased mitochondrial reactive oxygen species (ROS) (17); *osm-5* worms, which have reduced chemosensation (14) and *glp-1* worms, which are a model of germ line ablation (13). Since *glp-1* worms need to be grown at 25°C for the temperature-sensitive mutation to induce sterility and extend lifespan, a separate wild-type control grown at 25°C was used in all experiments.

We confirmed that all of the long-lived mutant strains have increased lifespan (**Fig. 1A; Fig. S1**). Importantly, by simultaneously measuring the lifespans of these strains in the same assay, the magnitude of lifespan extension can be directly compared between the long-lived mutants. In order of smallest to largest degree of lifespan extension were *ife-2* worms (26.3%), *clk-1* worms (33.4%), *sod-2* worms (37.2%), *eat-2* worms (45.6%), *osm-5* worms (65.4%), *nuo-6* worms (79.2%), *isp-1* (83.8%), *glp-1* worms (89.2%), and *daf-2* worms (138.4%). For the remaining figures, these strains will be arranged in order of the magnitude of lifespan extension, to easily visualize the extent to which stress resistance and expression of stress response genes correlates with lifespan.

**Figure 1.**
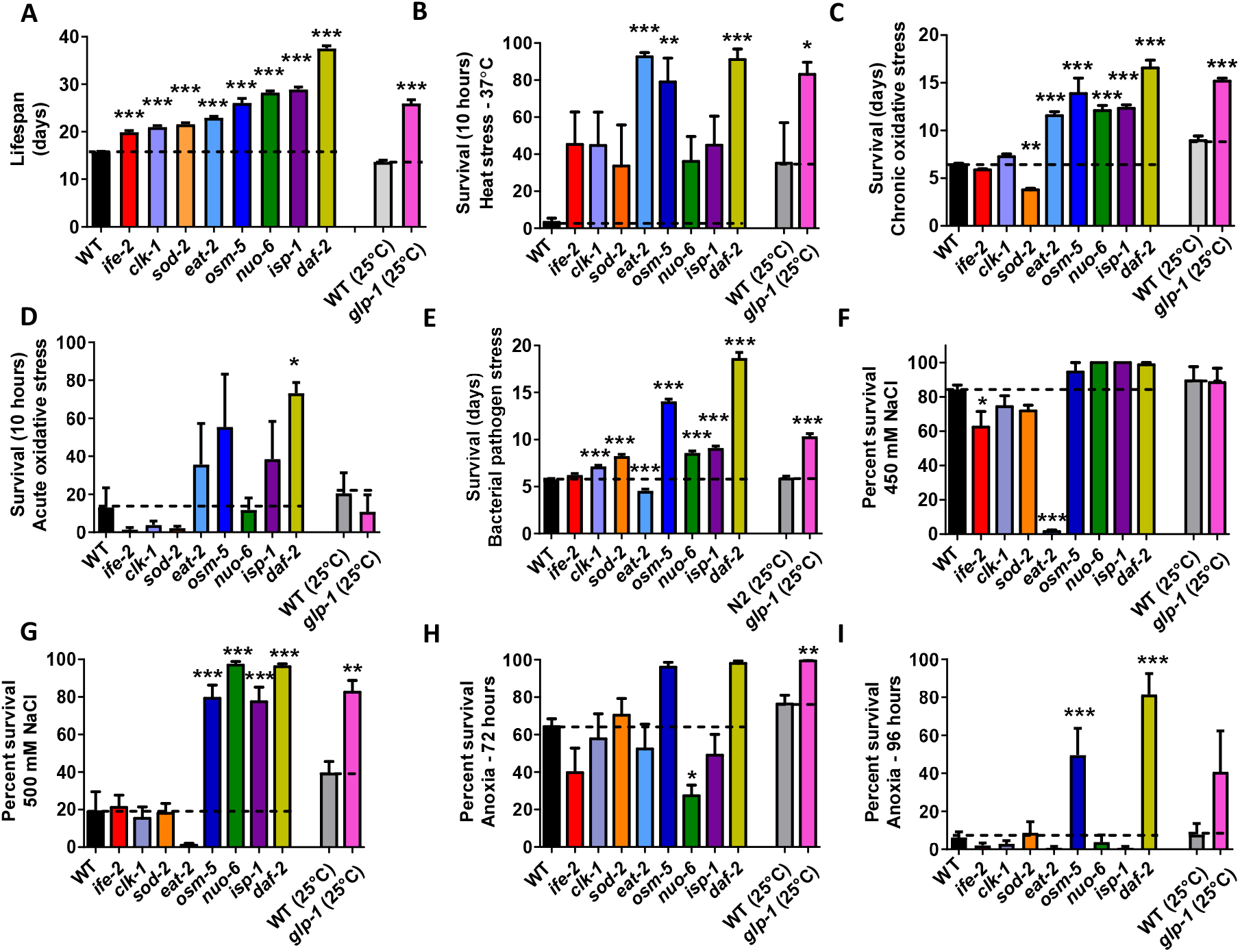
Long-lived mutants have increased resistance to multiple external stressors. **A**. The lifespans of nine different long-lived mutants were compared directly in the same assay. There was a range in the magnitude of lifespan extension observed across the nine mutants (for complete lifespan plots see **Figure S1**). **B**. All of the long-lived mutants exhibit increased resistance to heat stress at 37°C (for full time course see **Figure S2**). **C**. Six of the nine long-lived mutants have increased resistance to chronic oxidative stress (4 mM paraquat), while *sod-2* mutants have decreased resistance (for full time course see **Figure S3**). **D**. Four of the long-lived mutants show increased resistance to acute oxidative stress (300 μM juglone), while four mutants have decreased resistance (for full time course see **Figure S4**). **E**. All of the long-lived mutants except for *eat-2* mutants exhibit increased resistance to bacterial pathogen stress (for full time course see **Figure S5**). Resistance to osmotic stress was determined by exposing worms to 450 or 500 mM NaCl for 48 hours. **F**. At 450 mM NaCl, *ife-2* and *eat2* mutants show less resistance to osmotic stress compared to wildtype worms. **G**. At 500 mM NaCl, *osm-5, nuo-6, isp-1, daf-2*, and *glp-1* show increased resistance to osmotic stress. Resistance to anoxia was measured after 72 and 96 hours of anoxia. **H**. After 72 hours of anoxia, *nuo-6* mutants show decreased resistance to anoxia compared to wild-type worms, while *glp-1* shows increased resistance compared wild-type controls. **I**. After 96 hours of anoxia, *osm-5* and *daf-2* worms show increased resistance to anoxia compared to wild-type worms. For *glp-1* worms and their wild-type controls, worms were grown at 25°C during development and were shifted to 20 °C at adulthood. Error bars represent SEM. * p < 0.05, ** p < 0.01, *** p < 0.001. A minimum of three biological replicates were performed. Statistical analysis was performed using a one-way ANOVA with Dunnett’s multiple comparison test, except for *glp-1* which was compared to its wild-type control using a Student’s t-test.

### All long-lived mutants have increased resistance to one or more external stressor

To identify which types of stress resistance are most strongly correlated with longevity, we next quantified the relative resistance of the nine long-lived mutant strains to six external stressors using well-established stress paradigms including: heat stress (37°C), chronic oxidative stress (4 mM paraquat), acute oxidative stress (300 M juglone), bacterial pathogens (*P. aeruginosa* strain PA14), osmotic stress (450 or 500 mM NaCl), and anoxia (72 or 96 hours). Comparing the stress resistance of these mutants in the same assay also enabled us to identify genetic pathways driving the increased stress resistance in these long-lived mutants.

We found that all of the long-lived mutants showed increased resistance to heat stress compared to the wild-type worms, with *eat-2, osm-5, daf-2* and *glp-1* being the most resistant (**Fig. S2**; 10-hour time point is shown in **Fig. 1B**). The majority of the long-lived mutants also exhibited increased resistance to chronic oxidative stress, except for *ife-2* and *sod-2* worms, which have significantly decreased resistance (**Fig. S3**; average survival is shown in **Fig. 1C**). In contrast, only *eat-2, osm-5, isp-1* and *daf-2* worms have increased resistance to acute oxidative stress, while *ife-2, clk-1, sod-2* and *glp-1* worms are more sensitive (**Fig. S4**; 10-hour time point is shown in **Fig. 1D**). After exposure to bacterial pathogens, all of the long-lived mutants except for *eat-2* worms showed increased survival, with *osm-5* and *daf-2* worms having the greatest survival (**Fig. S5**; average survival is shown in **Fig. 1E**). In the osmotic stress assay, *ife-2* and *eat-2* show decreased resistance (**Fig. 1F**), while *osm-5, nuo-6, isp-1, daf-2* and *glp-1* have increased resistance (**Fig. 1G**). Finally, under conditions of anoxia, *nuo-6* worms have decreased survival while *glp-1, osm-5* and *daf-2* worms have increased survival compared to wild-type worms (**Fig. 1H, I**).

Overall, all of the long-lived mutants are resistant to at least one type of stressor (**Table S1**). Some of the long-lived mutants (*ife-2, sod-2, eat-2, nuo-6*) also show decreased resistance to certain external stressors. *daf-2* mutants are resistant to all six stressors tested and, in most assays, exhibited the greatest survival.

### Resistance to multiple different external stressors is significantly correlated with lifespan

To determine which types of stress resistance are correlated with lifespan, we compared the survival after exposure to each of the examined stressors to the average lifespan of the mutant. All six stressors had a significant, positive correlation between survival under stress and longevity with R^2^ values ranging from 0.3791 to 0.6630 (**Fig. 2A-F**). These results are consistent with a previous study which found a positive correlation between heat stress resistance and lifespan (R^2^ = 0.36), and UV resistance and lifespan (R^2^ = 0.42) (29). Resistance to bacterial pathogens (**Fig. 2D**, R^2^ = 0.6330) and resistance to chronic oxidative damage (**Fig. 2B**, R^2^ = 0.5628) exhibited the greatest correlation with lifespan, which may at least partially be due to the chronic nature of these assays.

**Figure 2.**
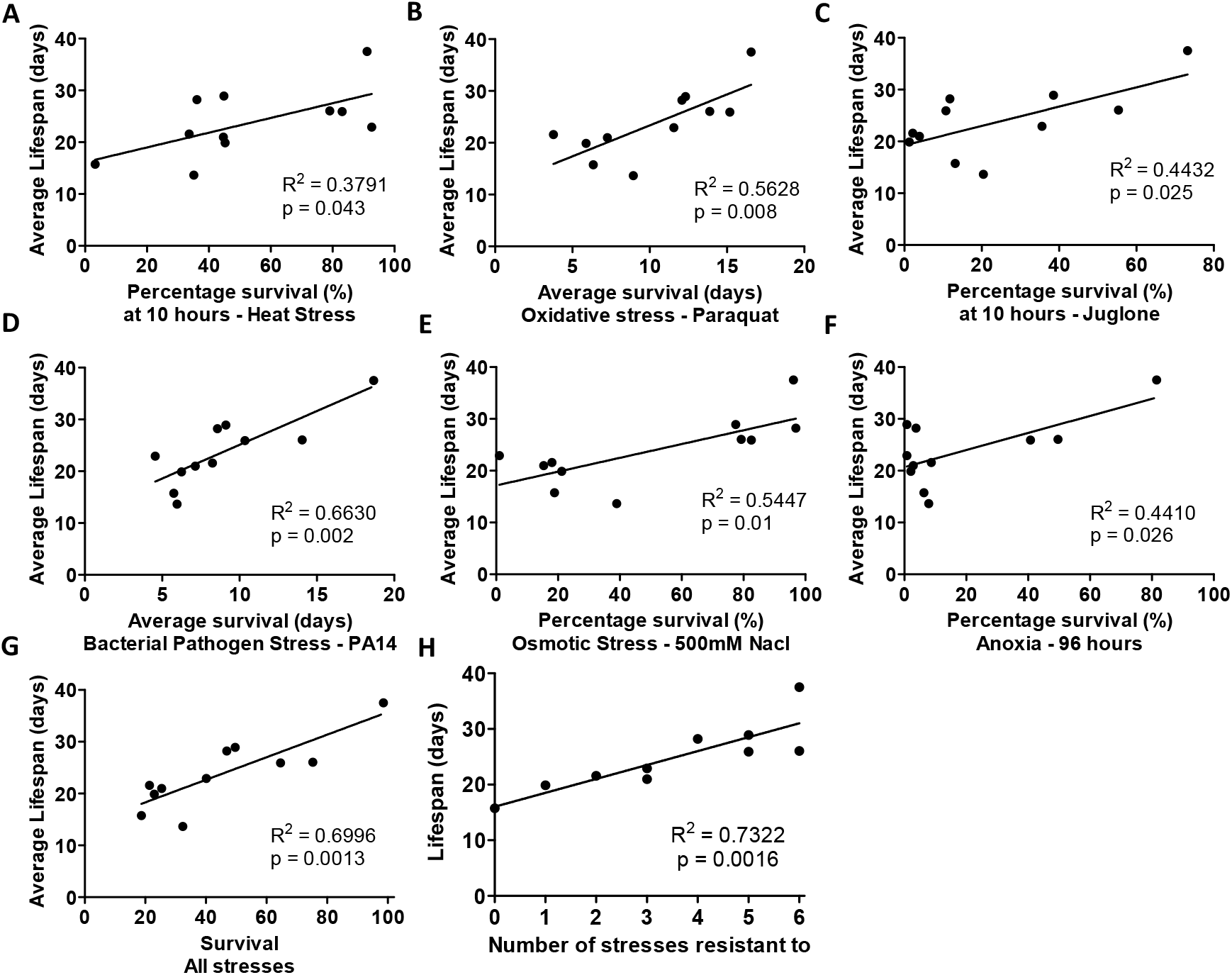
Resistance to multiple types of stress is significantly correlated with longevity. To determine the extent to which each type of stress resistance is correlated with lifespan, we compared the magnitude of lifespan extension to the percentage or duration of survival following exposure to exogenous stressors. There is a significant correlation with each of the six different types of stress resistance and lifespan including heat stress (**A**), chronic oxidative stress (**B**), acute oxidative stress (**C**), bacterial pathogens (**D**), osmotic stress (**E**) and anoxia (**F**). The highest degree of correlation was observed for bacterial pathogen stress. Combining all six types of stress resistance together into a combined survival score only slightly increased the degree of correlation with lifespan (**G**). The highest correlation with lifespan was achieved by using the number of stresses that a particular mutant is resistant to (**H**). * p < 0.05, ** p < 0.01, *** p < 0.001.

To evaluate whether resistance to multiple types of stressors is more predictive of lifespan than resistance to individual stressors, we combined the stress resistance data using two different approaches. First, we summed the relative stress resistance scores by setting the maximum stress resistance for each assay to 100% and expressed the stress resistance of each strain as a percentage of this maximum. This allowed us to combine the results from the six stress resistance assays at equal weight to generate an overall stress survival score. This combined stress survival score had only a marginally higher R^2^ value compared to the highest single stress R^2^ value (**Fig. 2G**, R^2^ = 0.6996). Second, we counted the numbers of stressors for which a strain showed significantly increased survival. Combining the stress resistance data in this way resulted in a slightly higher R^2^ value of 0.7322 (**Fig. 2H**). Overall, these results indicate that stress resistance is positively correlated with lifespan.

### Long-lived mutants exhibit upregulation of genetic targets of multiple stress response pathways

Having shown that all nine long-lived mutants that we tested display increased resistance to stress, we sought to explore the underlying mechanisms leading to stress resistance. We hypothesized that these mutants upregulate one or more stress response pathways. To evaluate the activation of each stress response pathway, we compared differentially expressed genes in the long-lived mutant strains to the target genes of established stress response pathways including the DAF-16-mediated stress response pathway (**Fig. 3A**; (33)), the p38-mediated innate immunity pathway (**Fig. 3B**; (34)), the HIF-1-mediated hypoxia response pathway (**Fig. 3C**; (35)), the SKN-1-mediated oxidative stress response pathway (**Fig. 3D**; (36)), the mitochondrial unfolded protein response (mitoUPR) pathway (**Fig. 3E**; (37, 38)), the cytoplasmic unfolded protein response (Cyto-UPR) pathway (**Fig. 3F**; (39, 40)), the ER-mediated unfolded protein response (ER-UPR) pathway (**Fig. 3G**; (41)), and antioxidant gene expression (**Fig. 3H**; (25)). Lists of the target genes from each stress response pathway, and the associated references are provided in **Table S2**.

**Figure 3.**
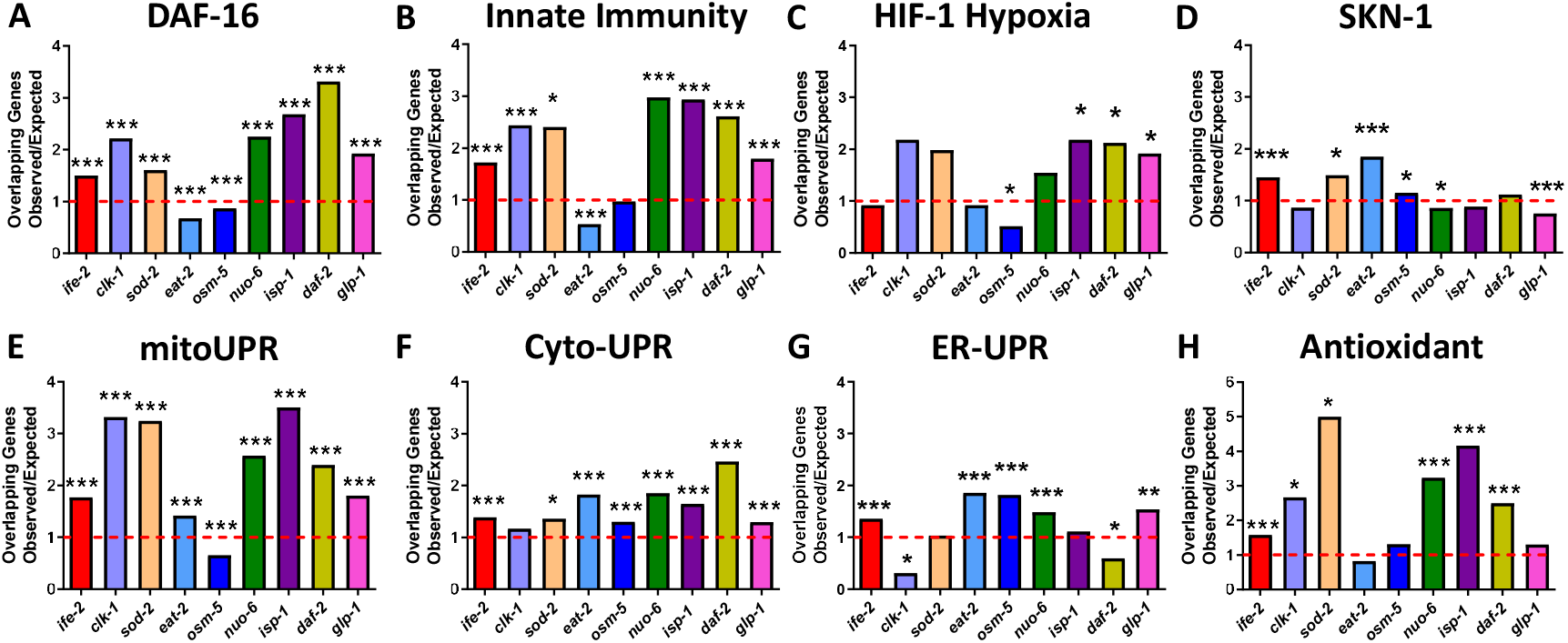
Long-lived mutants exhibit upregulation of genetic targets of multiple stress response pathways. Gene expression in the long-lived mutant strains was examined by RNA sequencing (RNA-seq) of six biological replicate per genotype of pre-fertile day 1 young adult worms. Differentially expressed genes that are significantly upregulated in the long-lived mutant strains were compared to genes that are upregulated by activation of different stress response pathways including the DAF-16-mediated stress response (**A**), the p38-regulated innate immune signaling pathway (**B**), the HIF-1-mediated hypoxia response (**C**), the SKN-1-mediated stress response (**D**), the mitochondrial unfolded protein response (mitoUPR) (**E**), the cytoplasmic unfolded protein response (Cyto-UPR) (**F**), the ER-mediated unfolded protein response (ER-UPR) (**G**), and antioxidant gene expression (**H**). Each of the stress response pathways is upregulated in multiple long-lived mutants. All of the long-lived mutants show a significant enrichment of genes involved in multiple stress response pathways. Venn diagrams comparing each individual long-lived mutant with each stress response pathway can be found in **Figures S6–S13**. * p < 0.05, ** p < 0.01, *** p < 0.001. p-value represents the significance of overlap between genes upregulated in the indicated long-lived mutant and genes upregulated by activation of the indicated stress response pathway.

In comparing the differentially expressed genes in each long-lived mutant to genes that are significantly modulated by activation by a specific stress response pathway, we determined the number of overlapping genes (see **Fig. S6–S13** for Venn diagrams), and then divided this number by the predicted number of overlapping genes expected if the genes were chosen at random to produce a ratio of observed/expected. All of the long-lived mutants had a significant enrichment of genetic targets from three or more stress response pathways (**Fig. 3**). In particular, we found that *ife-2* showed significant enrichment of genes from 7 stress response pathways; *sod-2, nuo-6, isp-1, daf-2* and *glp-1* showed enrichment of genes from 6 pathways; *clk-1* and *eat-2* showed enrichment of genes from 4 pathways; and *osm-5* showed enrichment of genes from 3 pathways. Interestingly, the most strongly enriched stress response pathway differed between strains (**Fig. S14**).

Amongst the stress response pathways, targets of the mitoUPR and the Cyto-UPR were enriched in eight long-lived mutants (**Fig. 3E,F**). Targets of the DAF-16-mediated stress pathway and the p38-mediated innate immunity pathway were enriched in seven mutants (**Fig. 3A,B**). Antioxidant genes were enriched in six mutants (**Fig. 3H**). Targets of the ER-UPR were enriched in five mutants (**Fig. 3G**). Targets of the SKN-1-mediated oxidative stress response were enriched in four mutants (**Fig. 3D**). Genes from the HIF-1-mediated hypoxia response were enriched in three mutants (**Fig. 3C**).

Finally, we examined the extent to which the ratio of observed/expected overlapping genes with each stress response pathway is correlated with longevity. Ratios for overlapping genes for both the DAF-16-mediated stress response pathway and the Cyto-UPR pathway have a significant correlation with lifespan (**Fig. S15**). These results suggest that the increased stress resistance in the long-lived mutants is caused by activation of multiple stress response pathways under unstressed conditions.

### Magnitude of upregulation of stress response genes in long-lived mutants

Having shown that there is a significant increase in the number of stress response genes showing differential expression in long-lived mutants, we wondered whether the magnitude of upregulation of these stress response genes impacts longevity. Accordingly, we measured the levels established target genes for each stress response pathway through qPCR: *sod-3* in the DAF-16 pathway (**Fig. 4A**; (26, 42)), *sysm-1/T24B8.5* in the innate immunity pathway (**Fig. 4B**; (43, 44)), *nhr-57* in the HIF-1 pathway (**Fig. 4C**; (26, 45)), *gst-4* in the SKN-1 pathway (**Fig. 4D**; (26, 46)), *hsp-6* in the mitoUPR pathway (**Fig. 4E**; (26, 47)), *hsp-16.2* in the Cyto-UPR pathway (**Fig. 4F**; (26, 48)), *hsp-4* in the ER-UPR pathway (**Fig. 4G**; (26, 49)), and the antioxidant gene *sod-5* (**Fig. 4H**; (50)). We observed a significant increase in the expression of *sod-3, sysm-1/T24B8.5, nhr-57, gst-4, hsp-6* and *sod-5*, in one or more long-lived mutants (**Fig. 4**). Moreover, the percentage increase in expression of either *sod-3* and *sod-5* was significantly correlated with longevity (R^2^ = 0.6697, 0.6312), while no correlation was observed with any of the other target genes (**Fig. S16**).

**Figure 4.**
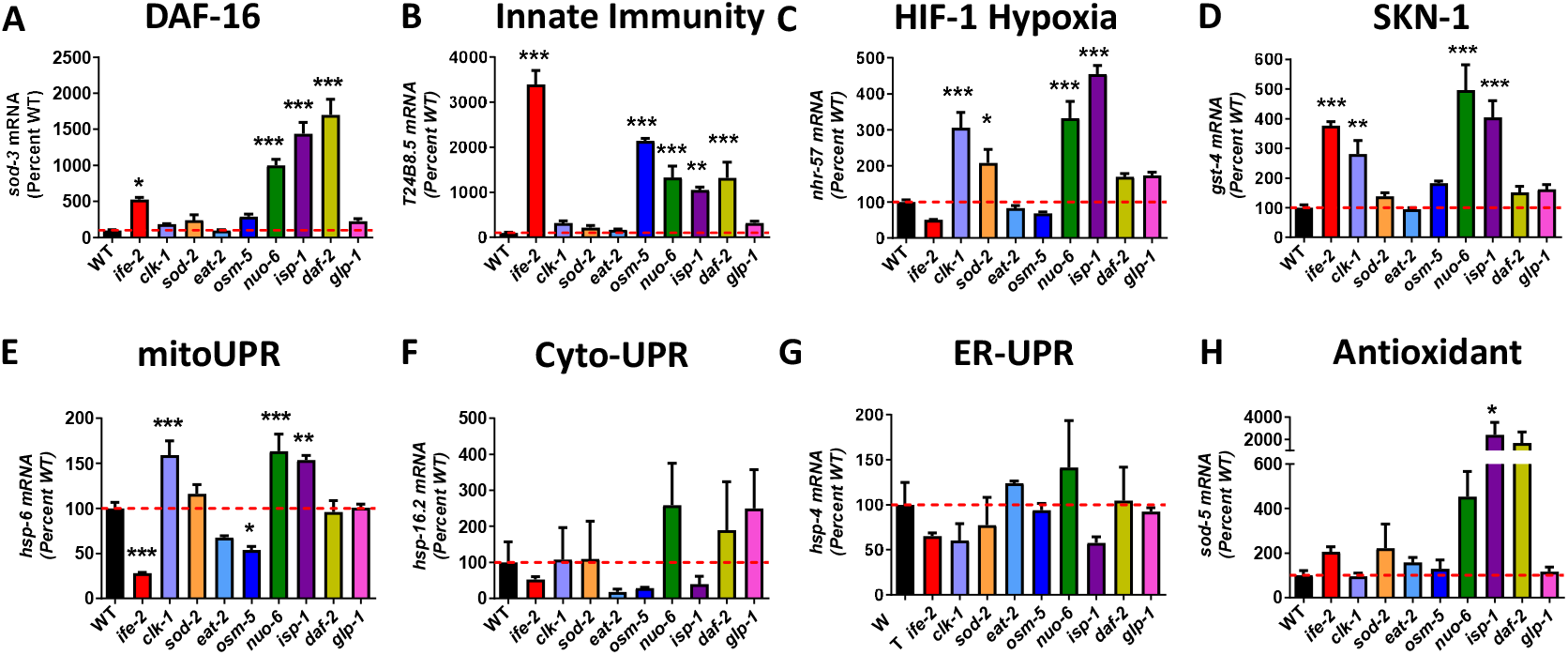
Magnitude of upregulation of stress response genes in long-lived mutants. Long-lived mutants show upregulation of genetic targets from each stress response pathway: *sod-3* (**A**) in the DAF-16-mediated stress response pathway; *sysm-1/T24B8.5* (**B**) in the p38-mediated innate immune pathway; *nhr-57* (**C**) in the HIF-1-mediated hypoxia pathway; *gst-4* (**D**) in the SKN-1-mediated oxidative stress response pathway; *hsp-6* (**E**) in the mitochondrial unfolded protein response (mitoUPR) pathway; *hsp-16.2* (**F**) in the cytoplasmic unfolded protein response (Cyto-UPR) pathway; *hsp-4* (**G**) in the endoplasmic reticulum unfolded protein response (ER-UPR) pathway; and *sod-5* (**H**) in the antioxidant pathway. Error bars represent SEM. * p < 0.05, ** p < 0.01, *** p < 0.001. Gene expression data is from RNA-sequencing of six biological replicates per strain. Statistical significance was assessed using a one-way ANOVA with Dunnett’s multiple comparison test.

### Genetic correlates of resistance to stress

To determine which individual genes are contributing to each type of stress resistance, we analyzed RNA-seq data from each long-lived mutant to identify genes whose expression is correlated with resistance to a specific stressor. We found that at least 71 genes (heat stress) and as many as 1115 genes (bacterial pathogens) exhibited a significant, positive correlation with each type of stress resistance, while 67 genes (heat stress) to 1014 genes (osmotic stress) exhibit a significant negative correlation with the different types of stress resistance (**Fig. S17, Table S3**). There were far more genes positively correlated with bacterial pathogen resistance than negatively correlated (7.4-fold more), similar to what we observed for lifespan. In contrast, resistance to osmotic stress had many more negatively correlated genes than positively correlated genes (2.2-fold more).

To determine the extent to which the same genes are driving different types of stress resistance, we compared genes that are both positively (**Fig. 5A**) and negatively (**Fig. 5B**) correlated with each type of stress resistance. We found that there are 20 genes correlated with resistance to five different stressors along with 69, 270, 666 and 2759 genes correlated with four, three, two, and one stressor, respectively (**Table S4**). For positively correlated genes, the greatest degree of overlap occurred between genes correlated with resistance to bacterial pathogens and genes correlated with resistance to anoxia (412 genes, 63% overlap), and between genes correlated with resistance to bacterial pathogens and genes correlated with resistance to acute oxidative stress (343 genes, 62% overlap) (**Fig. 5A**). For negatively correlated genes, the greatest overlap occurred between genes correlated with resistance to acute and chronic oxidative stress (100 genes, 36% overlap), and between genes correlated with resistance to osmotic stress resistance and genes correlated with bacterial pathogen resistance (73 genes, 49% overlap) (**Fig. 5B**). Combined, this suggests that similar genetic pathways can contribute to multiple types of stress resistance. In addition, there also exists some overlap in genes that are upregulated by activation of each stress response pathway (**Fig. S18** and (25)).

**Figure 5.**
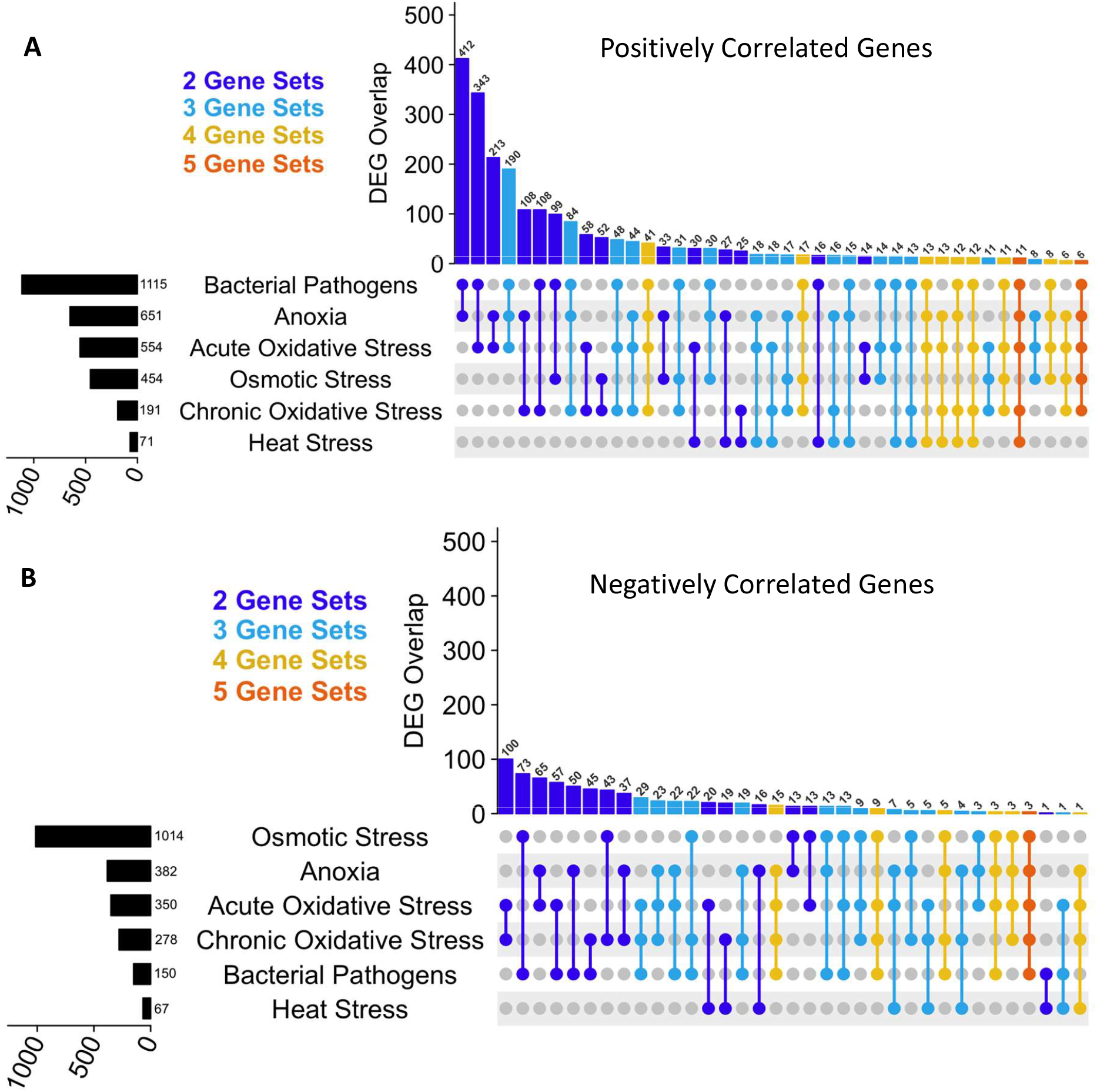
Genes correlated with resistance to different external stressors. The expression levels of all of the genes in the genome were determined by RNA sequencing in nine long-lived mutants. These expression levels were compared to the survival of these strains when exposed to different types of stress to determine which genes are correlated with stress resistance. **A**. An inclusive UpSetR plot comparing genes positively correlated with each type of stress resistance shows that there are many genes that are positively correlated with 2 or more types of stress resistance. **B**. An inclusive UpSetR plot comparing genes negatively correlated with each type of stress resistance shows that there are also multiple genes that are negatively correlated with 2 or more types of stress resistance. For panels A and B, the gene sets for each individual stressor are listed on the left and ordered by size from top to bottom. The height of each bar and the number above the bar indicate the number of genes in common between the gene sets indicated by the dots below the plot. The complete lists of genes that are significantly correlated with each type of stress resistance can be found in **Table S3**.

Resistance to each external stressor had a strong correlation to the expression of specific genes, with the maximum R^2^ values ranging from 0.71 for heat stress resistance to 0.91 for bacterial pathogen resistance (**Fig. 6**). This suggests that stress resistance is being determined by genetic factors. To determine if these same genetic factors also contribute to longevity, we examined whether the genes most strongly correlated with each type of stress resistance were also correlated with lifespan. We found that the genes most strongly correlated with chronic oxidative stress resistance and bacterial pathogen resistance were also correlated with lifespan (**Fig. 6**).

**Figure 6.**
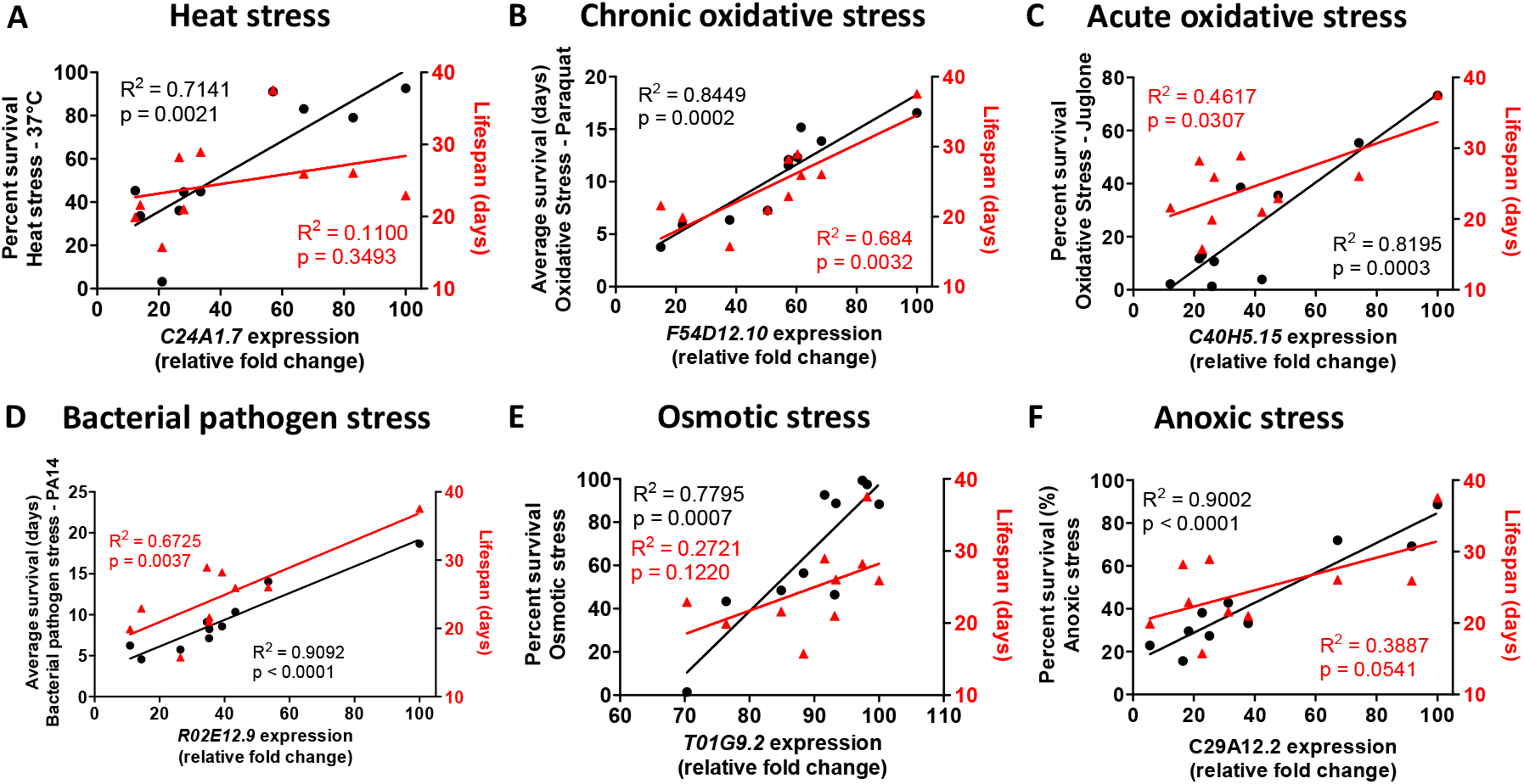
Expression levels of specific genes are highly correlated with resistance to stress. RNA sequencing data was used to determine which genes are most highly correlated with each type of stress resistance. The most highly correlated gene for each type of stress resistance are shown here. A full list of genes that are significantly correlated with each type of stress resistance can be found in **Table S3**. Of the genes that are most strongly correlated with stress resistance, genes correlated with chronic oxidative stress resistance (**B**) and genes correlated with bacterial pathogen resistance (**D**) were also correlated with lifespan. Stress resistance is indicated by black circles and black line (left Y-axis). Lifespan is indicated by red triangles and red line (right Y-axis). Relative fold change was calculated by dividing all of the expression values by the maximum expression value of all of the strains.

To understand the mechanisms by which genes correlated with stress resistance are acting to enhance resistance to stress, we looked for overrepresentation of these genes within Gene Ontology (GO) terms according to biological process, molecular function or cellular component (**Fig. S19**). There was an approximately three-fold enrichment within the biological processes “mitochondrial electron transport”, “aerobic respiration” and “regulation of response to oxidative stress” (**Fig. S19A**). This suggests that efficient generation of energy may help organisms respond to external stressors. For molecular functions, there was a 1.5-3 fold enrichment of “chitin binding”, structural constituent of cuticle”, “structural constituent of ribosome” and “oxidoreductase activity” (**Fig. S19B**). This suggests that enhancing the worm’s physical barrier to the environment promotes resistance to stress. Finally, for cellular components, there was a 1.5-3 fold enrichment of “mitochondrial respiratory chain complex I”, “cytosolic ribosomal subunit”, “collagen trimer” and “extracellular region” (**Fig. S19C**), again suggesting the importance of energy generation and physical barrier in enhancing resistance to external stressors.

### Genes contributing to stress resistance exhibit significant enrichment of genes contributing to longevity

To determine the extent to which the same genetic pathways are driving stress resistance and longevity on a broader scale, we compared genes that are correlated with each type of stress resistance to genes that are correlated with lifespan (Rudich et al., in preparation). We found that there was a significant degree of overlap between genes positively correlated with stress resistance and genes positively correlated with longevity ranging from 15% overlap (2.1-fold enrichment) to 84% overlap (11.5-fold enrichment), with the greatest degree of overlap occurring for resistance to bacterial pathogens (**Fig. 7; Fig. S20**). Genes that are negatively correlated with stress resistance were also negatively correlated with lifespan, though to a lesser extent than positively correlated genes (**Fig. S21**). These findings suggest that the same genes and genetic pathways are contributing to both stress resistance and lifespan.

**Figure 7.**
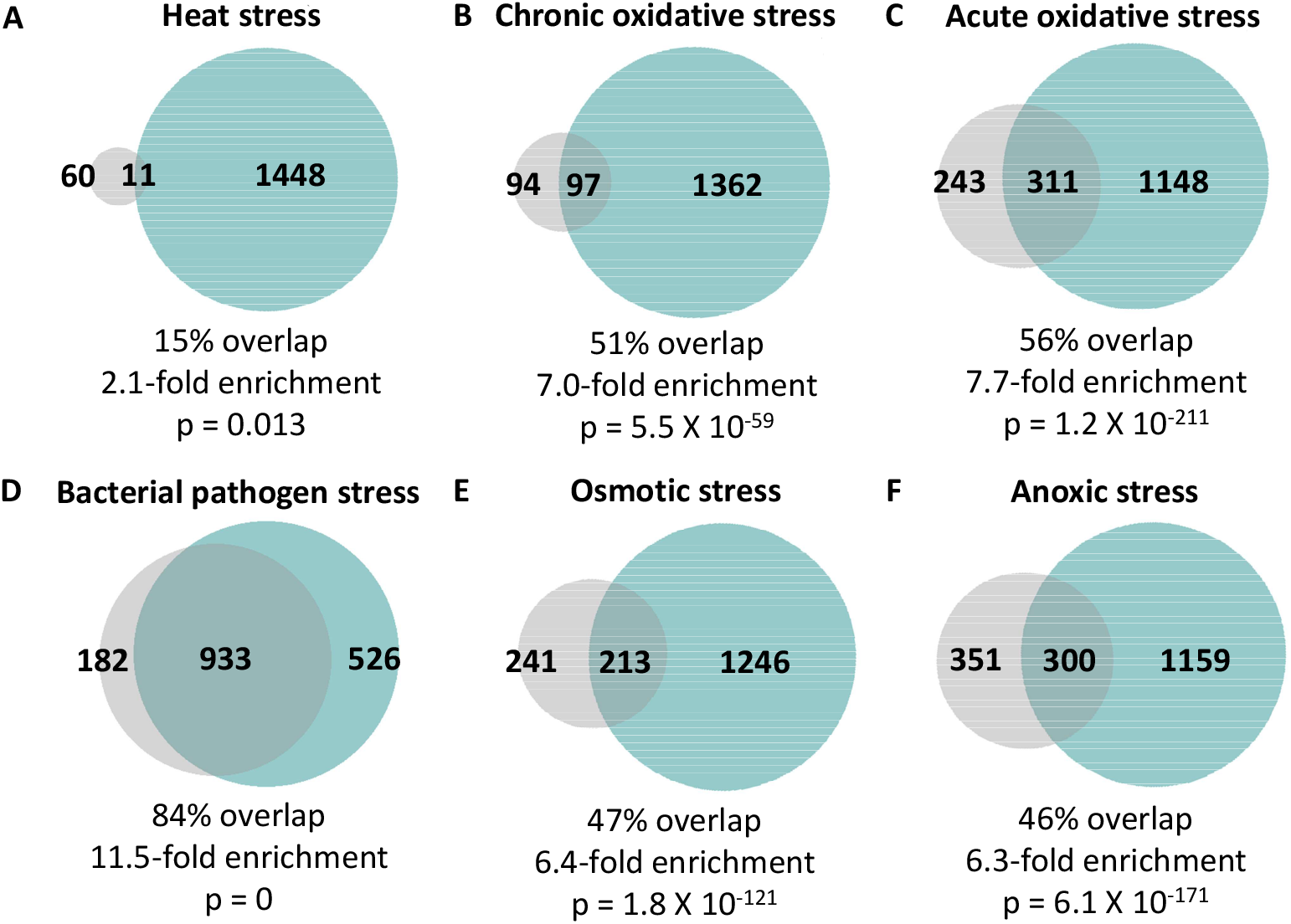
Highly significant overlap between genes the are correlated with resistance to stress and genes that are correlated with lifespan extension. In examining the degree of overlap between genes that are significantly correlated with lifespan (teal circle) and genes that are significantly correlated with resistance to stress (grey circle), we observed a significant degree of overlap with each of the six types of stress resistance that we examined including heat stress (**A**; 37°C), chronic oxidative stress (**B**; 4 mM paraquat), acute oxidative stress (**C**; 300 μM juglone), bacterial pathogen (**D**; *P. aeruginosa*), osmotic stress (**E**; 450-500 mM NaCl), and anoxic stress (**F**; 72-96 hours). The degree of enrichment ranged from 2.1 fold up to 11.5 fold with percent overlap between 15% and 84%. The highest degree of overlap with genes correlated with lifespan was with genes correlated with bacterial pathogen stress survival. The total number of genes that are unique to each circle (gene set) and number of overlapping genes are indicated. Percent overlap was calculated by dividing the number of overlapping genes by the number of genes positively correlated with each type of stress resistance. Enrichment was calculated as the number of overlapping genes observed divided by the number of overlapping genes expected by picking genes at random. The p-value indicates the significance of the difference between observed and expected numbers of overlapping genes.

### Disruption of stress response pathways has variable effects on longevity

To further examine the role of stress resistance in longevity, we reviewed previous data on the effect of transcription factors that regulate key stress response pathways on the lifespan of wild-type worms and long-lived mutants. Four pathways were consistently required for longevity. Disruption of the DAF-16-mediated stress response (*daf-16*) decreases lifespan in all of the long-lived mutants that we tested, and also decreases lifespan in wild-type worms (**Table 1**). A similar pattern was observed for the SKN-1-mediated oxidative stress response (skn-1), the cyto-UPR (hsf-1), and ER-UPR (*ire-1, xbp-1, pek-1, atf-6*), although these genes were only interrogated in a subset of the long-lived mutants that we studied for this report (**Table 1**). Thus, the DAF-16-mediated stress response, the SKN-1 mediated oxidative stress response, the cyto-UPR and the ER-UPR are required for normal lifespan and may be required for extended lifespan.

**Table 1.** Effect of disrupting transcription factors controlling stress response pathways on lifespan in wild-type worms and long-lived mutants. The left most column indicates the genotype of the strain. Each other column represents the stress response pathway indicated in bold in the top row. DAF-16 = DAF-16-mediated stress response; Innate Immunity = p38-regulated innate immune signaling pathway; HIF-1 = HIF-1-mediated hypoxia response; SKN-1 = SKN-1-mediated stress response; mitoUPR = mitochondrial unfolded protein response; Cyto-UPR = the cytoplasmic unfolded protein response; ER-UPR = ER-mediated unfolded protein response; and Antioxidant = antioxidant genes. “↑” disruption of the transcription factor increased lifespan; “↓” disruption of the transcription factor decreased lifespan; “=” disruption of the transcription factor did not affect lifespan. NR = not reported

|  | **DAF-16** | **Innate Immunity** | **HIF-1** | **SKN-1** | **mitoUPR** | **Cyto-UPR** | **ER-UPR** | **Antioxidant** |
| --- | --- | --- | --- | --- | --- | --- | --- | --- |
| **WT** | *daf-16* ↓(4, 22, 72, 73) | *nsy-1* =(44, 74)<br>↓(43, 74)<br>*sek-1* =(44, 74)<br>↓(43)<br>*pmk-1* =(43, 44, 74) | *hif-1* ↑(75)<br>↓(76) | *skn-1* ↓(38, 77) | *atfs-1* ↑(78)<br>= (56, 58)<br>*ubl-5* =(79)<br>*dve-1* ↓(79) | *hsf-1* ↓(22, 80-82) | *ire-1* ↓(83, 84)<br>*xbp-1* ↓(83)<br>*pek-1* ↓(83)<br>*atf-6* ↓(83) | *sod-1* ↓(17, 85)<br>*sod-2* ↑(17, 85)<br>*sod-3* =(17, 85)<br>*sod-4* =(17, 85)<br>*sod-5* =(17, 85) |
| ***ife-2*** | *daf-16* ↓(15, 16) | NR | NR | *skn-1* ↓(86) | NR | *hsf-1* ↓(87) | NR | NR |
| ***clk-1*** | *daf-16* ↓(61) | *pmk-1* ↓(88) | *hif-1* ↓(76) | *skn-1* ↓(89) | *atfs-1* ↓(56)<br>*ubl-5* ↓(79) | NR | NR | *sod-1* ↓(54)<br>*sod-2* ↑(17, 54)<br>*sod-3* =(54)<br>*sod-4* =(54)<br>*sod-5* ↓(54) |
| ***sod-2*** | *daf-16* ↓(61) | *pmk-1* ↓(88) | NR | *skn-1* ↓(88) | *atfs-1* =(56) | NR | NR | *sod-1* ↑(17)<br>*sod-3* ↓(17)<br>*sod-4* =(17)<br>*sod-5* ↓(17) |
| ***eat-2*** | *daf-16* ↓(12) | *sek-1* ↓(43) | *hif-1* =(76, 90) | *skn-1* ↓(89) | *ubl-5* =(79)<br>*dve-1* ↓(79) | *hsf-1* ↓(80) | *ire-1* ↓(83) | *sod-2* ↑(17) |
| ***osm-5*** | *daf-16* ↓(14) | NR | *hif-1* =(76) | NR | NR | NR | NR | NR |
| ***nuo-6*** | *daf-16* ↓(61) | *nsy-1* ↓(44)<br>*sek-1* ↓(44)<br>*pmk-1* ↓(44)<br>*atf-7* ↓(44) | *hif-1* ↓(91) | *skn-1* ↓(88) | *atfs-1* ↓(56) | *hsf-1* ↓(88) | NR | NR |
| ***isp-1*** | *daf-16* ↓(61) | *nsy-1* ↓(44)<br>*sek-1* ↓(44)<br>*pmk-1* ↓(44)<br>*atf-7* ↓(44) | *hif-1* ↓(76) | *skn-1* ↓(88) | *atfs-1* ↓(56)<br>*ubl-5* ↓(79)<br>*dve-1* ↓(79) | *hsf-1* ↓(80)<br>*hsf-1* ↓(88) | NR | *sod-1* ↓(88)<br>*sod-2* ↓(17)<br>*sod-3* ↓(55)<br>*sod-4* ↓(88)<br>*sod-5* ↓(55) |
| ***glp-1*** | *daf-16* ↓(92, 93) | NR | *hif-1* =(76) | *skn-1* ↓(94) | *atfs-1* ↓(95) | *hsf-1* ↓(96) | NR | *sod-2* ↑(17) |
| ***daf-2*** | *daf-16* ↓(4, 22) | *sek-1* ↓(43, 97)<br>*pmk-1* ↓(22, 97, 98)<br>*atf-7* ↓(43) | *hif-1* ↑(75)<br>= (76, 99) | *skn-1* ↓(38, 77) | *ubl-5* =(79)<br>*dve-1* ↓(79) | *hsf-1* ↓(22, 80) | *ire-1* ↓(83)<br>*xbp-1* ↓(83)<br>*pek-1* ↓(83)<br>*atf-6* ↓(83) | *sod-2* ↓(100)<br>=(17)<br>*sod-3* ↑(100)<br>*sod-4* ↑(100)<br>*sod-12345* ↓(22) |

For innate immunity, disruption of *pmk-1* specifically reduced the lifespan of long-lived mutants (*clk-1, sod-2, nuo-6, isp-1, daf-2*) but had no effect on wild-type worms (**Table 1**), suggesting that this pathway contributes to the enhanced longevity of long-lived mutants but is not needed for wild-type lifespan. Meanwhile, the HIF-1-hypoxia pathway appears to be specifically required for the longevity of long-lived mitochondrial mutants, as disruption of *hif-1* decreased lifespan in *clk-1, isp-1* and *nuo-6* worms but not in other long-lived mutants (*eat-2, osm-5, glp-1, daf-2*) (**Table 1**). Similarly, the mitoUPR is primarily required for extended lifespan in mitochondrial mutants, as disruption of *ubl-5* decreases *clk-1* and *isp-1* lifespan, but not *eat-2* or *daf-2* (**Table 1**). Finally, disruption of superoxide dismutase antioxidant genes has variable effects on lifespan depending on the subcellular localization of the SOD protein, and the strain being examined (**Table 1**).

These results show that each of the stress response pathways we examined here are required for the extended longevity of at least some of the long-lived mutants studied. In addition, many of these stress response pathways are also required for wild-type lifespan. Combined, this emphasizes the importance of stress response signaling pathways for longevity.

## DISCUSSION

The survival of an organism depends upon its ability to resist physiological stress. Animals not only have to survive exogenous stressors in their environment such as changes in temperature, changes in oxygen availability, and presence of pathogens, but also internal stressors produced as by-products of cellular processes. The activation of stress pathways in response to physiological stressors is important to maintain cellular homeostasis amidst fluctuations in the environment and evolutionarily conserved. By quantifying resistance to multiple external stressors in long-lived *C. elegans* mutants and analyzing their gene expression, we found that long-lived mutants have increased resistance to stress due to upregulation of multiple stress response pathways. Genes correlated with stress resistance exhibit a highly significant overlap with genes correlated with lifespan suggesting that similar genetic pathways contribute to both phenotypes.

### Increased resistance to stress is associated with extended longevity

Since the discovery of genetic mutant strains that have extended lifespan, multiple groups have examined their ability to withstand different external stressors (51). *daf-2* mutants are the most well characterized in this regard, and have increased resistance to oxidative stress, heat stress, osmotic stress, anoxia, heavy metals, and bacterial pathogens (18–22). Similarly, *eat-2* mutants have increased resistance to heat stress (16); *ife-2* mutants have increased resistance to heat stress and oxidative stress (15, 16); *glp-1* mutants have increased resistance to bacterial pathogens, heat stress and oxidative stress (52, 53); *clk-1* mutants have increased resistance to chronic oxidative stress (54); *isp-1* mutants have increased resistance to heat stress, oxidative stress, osmotic stress and bacterial pathogens (44, 55); and *nuo-6* worms have increased resistance to heat stress, oxidative stress, osmotic stress and bacterial pathogens (44, 56).

In this study, we extended previous findings by comprehensively examining resistance to six of the most well-studied stress paradigms in nine long-lived mutants representing multiple different pathways of lifespan extension (**Fig. 1**). Our results show that all of the examined long-lived mutants have increased resistance to at least one external stressor, with some mutants having increased resistance to all examined stressors (*daf-2, osm-5*), while other mutants are resistant to only one (*ife-2*). The number of stressors that a mutant is resistant to is correlated with the magnitude of their lifespan extension, suggesting that resistance to a wide array of stressors is beneficial for longevity, even under laboratory conditions, which are believed to be relatively unstressful. An alternative explanation for the correlation between stress resistance and lifespan, is that the same genetic pathways contribute to both aging and stress resistance, which is supported by our gene expression studies.

All of the types of stress resistance that we examined showed a significant positive correlation with lifespan, with resistance to bacterial pathogens having the strongest relationship with lifespan (**Fig. 2**). This extends the findings of a previous study that showed a positive correlation between lifespan and thermotolerance, UV resistance and juglone resistance using different sets of mutants (29). In our study, all of the long-lived mutants showed increased resistance to heat stress, suggesting that this type of stress resistance may be more important for longevity than other types. However, others have observed decreased resistance to heat stress in long-lived *daf-28* mutants (29, 57). Thus, for each of the external stressors we examined, there are one or more long-lived mutants that exhibit decreased survival. This indicates that increasing lifespan is not sufficient to provide protection against any specific external stressors.

### Stress resistance can be experimentally dissociated from lifespan

To untangle the relationship between stress resistance and longevity, researchers have experimentally manipulated stress resistance pathways and examined the resulting effects on lifespan. While disrupting stress response pathways typically decreases lifespan in both wild-type worms and long-lived mutants (see **Table 1** and references therein), there are multiple exceptions.

Mutations can decrease resistance to one or multiple stressors, but have no effect on lifespan, or even increase lifespan. Disruption of the mitoUPR transcription factor ATFS-1 decreases resistance to multiple stressors (56), but is reported to either have no effect on lifespan (56, 58) or increase lifespan (24). Deletion of *sod-2* decreases resistance to oxidative stress, but increases lifespan (17). Similarly, disruption of *clk-1* decreases resistance to acute oxidative stress but increases lifespan (54). Here we show that *ife-2* mutants have decreased resistance to oxidative stress and osmotic stress, *eat-2* mutants have decreased resistance to osmotic stress and bacterial pathogens, and *glp-1* worms have decreased resistance to acute oxidative stress, but all three mutants still show increased lifespan (**Table S1**). Worms with mutations disrupting all five superoxide dismutase genes (*sod*) have decreased resistance to multiple stressors, but do not have decreased lifespan (22, 59). Disruption of both glycerol-3-phosphate dehydrogenase (*gpdh*) genes or deletion of the putative E3 ubiquitin ligase gene *nhl-1* makes *daf-2* worms more sensitive to osmotic stress, but increases their lifespan (22). Disruption of the p38-mediated innate immunity pathway through deletion of *pmk-1* decreases resistance to bacterial pathogens but does not affect lifespan (22). Combined, these examples demonstrate that decreasing resistance to stress is not sufficient to decrease lifespan.

There are also examples in which increasing stress resistance does not increase lifespan. Mutations in *daf-4* or *daf-7* result in increased resistance to heat stress, but do not extend longevity (20). Constitutive activation of ATFS-1 increases resistance to multiple stressors but decreases lifespan (25). Disruption of the transcription elongation regulator TCER-1 increases resistance to bacterial pathogens, heat stress and oxidative stress in *glp-1* mutants, but decreases their lifespan (53, 60). Thus, increasing stress resistance is not sufficient to increase lifespan. Combined, these examples indicate that stress resistance can be experimentally dissociated from longevity.

### Long lived mutants exhibit upregulation of multiple stress response pathways

To better understand the molecular mechanisms underlying the increased stress resistance in the long-lived mutants, we compared gene expression in these mutants to genes modulated by activation of established stress response pathways. We found that all nine of the long-lived mutants exhibit a statistically significant enrichment of genes that are upregulated by activation of multiple stress response pathways. The greatest number of mutants showed enrichment of genetic targets of the mitoUPR pathway (8 of 9), the cyto-UPR pathway (8 of 9), the DAF-16-mediated stress response pathway (7 of 9) and the p38-mediated innate immunity pathway (7 of 9), suggesting that these stress response pathways may be more important contributors to stress resistance and longevity of these mutants. These findings extend our previous work in which we observed a significant upregulation of DAF-16 target genes (61), mitoUPR target genes (56), innate immunity genes (44) and antioxidant genes (54, 55) in the long-lived mitochondrial mutants *clk-1, isp-1*, and *nuo-6*.

While the specific contribution of each of these stress response pathways to stress resistance in long-lived mutant strains has not been examined in most cases, we and other groups have examined the effects of stress response pathways on the longevity of long-lived mutant strains (summarized in **Table 1**). In many cases, including DAF-16, SKN-1, DVE-1, HSF-1, IRE-1, XBP-1, PEK-1, ATF-6 and in some reports NSY-1, SEK-1, and HIF-1, disruption of the transcription factor regulating the stress response pathway decreases lifespan in wild-type worms and long-lived mutants. As a result, the contribution of the stress response pathway to the extended longevity of the long-lived mutant is difficult to assess because the decrease in wild-type lifespan suggests a generally detrimental effect on longevity. The clearest contributions to longevity come from stress response pathways that do not impact the lifespan of wild-type animals including the p38-mediated innate immune signaling pathway and the mitoUPR. Deletion of the p38-mediated innate immune signaling pathway regulator, *pmk-1*, decreases the lifespan of *sod-2, clk-1, isp-1, nuo-6* and *daf-2* mutants, while having no effect on wild-type lifespan (**Table 1**). Disrupting the main regulators of the mitoUPR pathway, *atfs-1* or *ubl-5*, decreases the lifespan of *clk-1, nuo-6, isp-1* and *glp-1* mutants, but has no effect on wild-type worms (**Table 1**). Thus, both the p38-mediated innate immune signaling pathway and the mitoUPR contribute to the longevity of multiple long-lived mutants, while the role of other stress response pathways is less clear because disruption of these pathways decreases wild-type lifespan.

### The same genetic pathways affect stress resistance and longevity

From our results here and previous studies, the significant correlation between stress resistance and lifespan cannot be explained by models in which increased stress resistance causes extended longevity or vice versa, as multiple counterexamples exist. To gain insight into this relationship, we examined the extent to which the genes driving stress resistance overlap with genes driving longevity. For each of the six types of stress resistance examined, there was a statistically significant overlap between genes correlated with stress resistance and genes correlated with longevity, suggesting that many of the same genes that contribute to stress resistance also contribute to longevity. Based on the high degree of overlap between genes controlling stress resistance and genes controlling longevity, we propose that the significant correlation between stress resistance and aging is due to a large group of genes that contribute to both stress resistance and longevity (**Fig. S22**). Whenever one of these overlapping genes is modulated, both phenotypes are affected. There are also sets of genes that contribute only to lifespan or only to stress resistance. Modulation of these genes can affect lifespan independently of stress resistance, or vice versa, thereby allowing for the experimental dissociation of stress resistance and lifespan.

### Innate immunity and longevity

Aging is associated with dysregulation of the immune system (62). In *C. elegans*, aging results in increased susceptibility to bacterial infections as well as a decline in activation of the p38-mediated innate immunity pathway (26, 63), which is the main pathway for pathogen defense (64). Of all of the types of stress resistance that we examined, we observed the strongest relationship between resistance to bacterial pathogens and longevity. Lifespan was most highly correlated with resistance to bacterial pathogens (**Fig. 2**), and genes correlated with lifespan had the highest overlap with genes correlated with resistance to bacterial pathogens (**Fig. 7**; 84%). In addition, seven of the nine long-lived mutants we examined exhibited a significant upregulation of target genes of the p38-mediated innate immune signaling pathway (**Fig. 3**). Combined, these results indicate that activation of pathways enabling survival of bacterial pathogens exposure promotes longevity. The importance of innate immunity for longevity is supported by recent work from our laboratory and others showing that the p38-mediated innate immune signaling pathway is required for the longevity of multiple long-lived mutants including *clk-1, sod-2, eat-2, nuo-6, isp-1*, and *daf-2* (see **Table 1**). This appears to be true even under unstressful laboratory conditions in the absence of bacterial pathogens. While the OP50 strain of *E. coli* bacteria, which is the standard food source for maintaining *C. elegans*, can have detrimental effects on lifespan due to proliferation in the pharynx and intestine (65, 66), the p38-mediated innate immune signaling pathway is still required for the extended lifespan of long-lived mutants when they are being fed non-proliferating bacteria (43, 44).

## CONCLUSION

Different pathways of lifespan extension, examined via long-lived *C. elegans* mutants, all induce enhanced resistance to at least one external stressor and significant upregulation of multiple stress response pathways. All six types of stress resistance examined are significantly correlated with longevity, with the strongest correlation being with resistance to bacterial pathogens. By identifying the genes that are most highly correlated with each type of stress resistance, we found that the same genetic pathways control both resistance to stress and longevity, and that the strongest relationship exists between resistance to bacterial pathogens and lifespan. This indicates that the correlation between stress resistance and lifespan is due to a large group of genes that control both phenotypes. Overall, this work demonstrates a role for stress response pathways in determining lifespan and emphasizes the importance of innate immune signaling for longevity.

## METHODS

### Strains and maintenance

All *C. elegans* strains were obtained from the Caenorhabditis Genetics Center (CGC): N2 (wild-type), *ife-2 (ok306), clk-1(qm30), sod-2(ok1030), eat-2(ad1116), osm-5(p813), nuo-6(qm200), isp-1(qm150), daf-2(e1370), glp-1(e2141)*. All strains were grown and maintained in nematode grown medium (NGM) plates at 20°C except *glp-1(e2141), which* was maintained at 20°C, but grown at 25°C to induce sterility and lifespan extension in experimental worms. Plates were seeded with OP50 *E. coli* as a food source.

### Lifespan

Lifespan experiments were conducted at a temperature of 20°C using at least 3 biological replicates with a minimum of 40 worms per strain for each trial. Survival was measured every day until death. Plates contained 25μM 5’-fluorodeoxyuridine (FUdR) to minimize internal hatching of progeny (67). Survival data was pooled across multiple trials. Worms with internal hatching or extruded vulva were censored.

### Stress Assays

All stress assays were conducted using at least 3 biological replicates with a minimum of 20 worms per plate at a temperature of 20°C.

### Oxidative stress

For acute oxidative stress, young adult worms were transferred onto freshly poured agar plates containing 300 μM juglone. Survival was measured every 2 hours for a total of 10 hours. For chronic oxidative stress, young adult worms were transferred onto agar plates containing 4 mM paraquat and 100 μM FUdR. Survival was measured daily until death.

### Heat stress

Resistance to heat stress was tested by transferring young adult worms to agar plates and placing plates in 37°C. Survival was measured every 2 hours for a total of 10 hours.

### Osmotic stress

Resistance to heat stress was tested by transferring young adult worms to agar plates containing 450 mM or 500 mM NaCl in the NGM plates. Survival was measured after 48 hours.

### Anoxic stress

Resistance to anoxic stress was tested by transferring young adult worms to agar plates and putting the plates in BD Bio-Bag Type A Environmental Chambers (Becton, Dickinson and Company, NJ). Survival was measured after 72 or 96 hours.

### Bacterial pathogen stress

Resistance to bacterial pathogen stress was performed using the slow-killing assay as previously described (43, 44). *P. aeruginosa* liquid culture were grown in Luria–Bertani (LB) media overnight and were used to seed NGM plates for the slow-killing assay. Plates were incubated at 37°C for 24 hours and then at 20°C for 24 hours. L4 worms were transferred to NGM plates containing 100mg/L FuDR and grown with OP50 until day 3 of adulthood. Adults were transferred to PA14-seeded plates containing 20mg/L FuDR. Assay was conducted at 20°C. Worms were checked daily until death.

### RNA sequencing and bioinformatic analysis

RNA sequencing was performed on young adult worms collected from six independent samples for each strain as described previously (55). RNA-seq data is available on NCBI GEO (GSE93724(61) GSE110984 (56), GSE179825) and was analyzed by the Harvard School of Public Health Bioinformatics core for this paper. For read mapping and expression level estimation, we used an RNA-seq pipeline from the bcbio-nextgen project (https://bcbio-nextgen.readthedocs.org/en/latest/) to process samples. We examined quality of the raw reads using FastQC (http://www.bioinformatics.babraham.ac.uk/projects/fastqc/) and trimmed any reads that contained contaminant sequences and low quality sequences with cutadapt (http://code.google.com/p/cutadapt/). We used STAR (68) to align trimmed reads to the Ensembl build WBcel235 (release 90) of the *C. elegans* genome. To check alignment quality, we checked for evenness of coverage, rRNA content, genomic context of alignments, and complexity. For expression quantification, we used Salmon (69) for identifying transcript-level abundance estimates and then used R Bioconductor package tximport (70) for collapsing down to the gene level. To validate sample clustering from the same batches across different mutants, we used principal components analysis (PCA) and hierarchical clustering methods. We used R Bioconductor package DESeq2 (71) to perform differential gene expression analysis. For each comparison between wildtype and mutant, a false discovery rate (FDR) threshold of 0.01 was set to identify significant genes. To adjust for batch effects, we included batch as a covariate in the linear model for datasets in experiments that were run across two batches.

### Overlapping genes

Lists of the differentially expressed genes from mutant were compared to genes modulated by each stress pathway. Comparisons were made between differentially expressed genes of the same direction of change. The hypergeometric test was used to compute significance of overlap. Venn diagrams were made from the online tool BioVenn (https://www.biovenn.nl/).

### Gene Ontology Analysis

PANTHER Classification System (version 16.0) was used for functional analysis of genes correlated with stress. Statistical overrepresentation analysis of Gene Ontology (GO) terms was performed by inputting the WormBase IDs. Fisher’s Exact test was used to calculate false discovery rates to determine the significantly enriched GO terms.

### Statistical analysis

Experiments were performed with the experimenter blinded to the genotypes of the worms being tested. A minimum of three biological replicates on different days were performed for each assay. All statistical analyses were performed using GraphPad Prism version 5.01. Survival plots for lifespan assays, oxidative stress assays, and bacterial pathogen stress assays were analyzed using a Log-rank test. Heat stress assays were analyzed using repeated measures ANOVA. Osmotic stress and anoxic stress assays were analyzed using one-way ANOVA with Dunnett’s multiple comparison test.

## Acknowledgments

We would like to thank the laboratory of Dr. Don Sheppard for providing *Pseudomonas aeruginosa* strain PA14. Some strains were provided by the CGC, which is funded by NIH Office of Research Infrastructure Programs (P30 OD010440). We would also like to acknowledge the *C. elegans* knockout consortium and the National Bioresource Project of Japan for providing strains used in this research.

## Competing interests

The authors have declared that no competing interests exist.

## Author Contributions

Conceptualization: JVR. Methodology: SKS, PR, MM, JVR. Investigation: SKS, PR, MM, JVR. Analysis: SKS, PR, MM, JVR. Visualization: SKS, PR, MM, JVR. Writing – original draft: SKS, JVR. Writing – review and editing: SKS, PR, JVR. Supervision: JVR.

## Materials & Correspondence

Correspondence and material requests should be addressed to Jeremy Van Raamsdonk.

## Data availability

RNA-seq data has been deposited on GEO: GSE93724, GSE110984 and GSE179825. All other data and strains generated in the current study are included with the manuscript or available from the corresponding author on request.

## Funding

This work was supported by the Canadian Institutes of Health Research (CIHR; http://www.cihr-irsc.gc.ca/; JVR), the Natural Sciences and Engineering Research Council of Canada (NSERC; https://www.nserc-crsng.gc.ca/index_eng.asp; JVR), and the National Institute of General Medical Sciences (NIGMS; https://www.nigms.nih.gov/; JVR) by grant number R01 GM121756. JVR is the recipient of a Senior Research Scholar career award from the Fonds de Recherche du Québec Santé (FRQS) and Parkinson Quebec. SKS is supported by a FRQS Doctoral Award. PDR is supported by a FRQS Postdoctoral award. The funders had no role in study design, data collection and analysis, decision to publish, or preparation of the manuscript.

## Supporting Information

**Figure S1.**
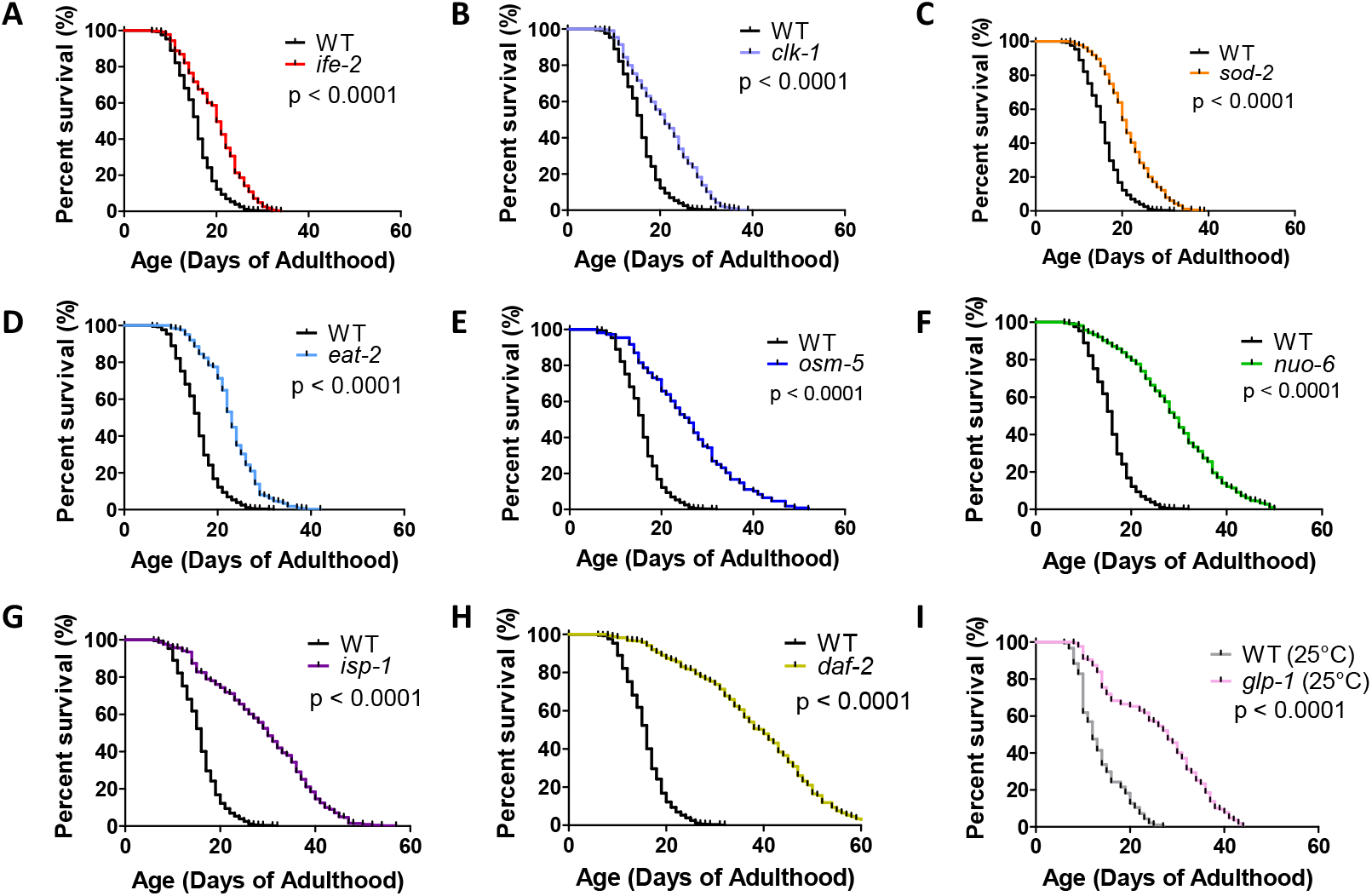
Long-lived mutants exhibit different magnitudes of lifespan extension. In order to compare the length of lifespan extension between different long-lived mutants, we measured the lifespan of nine long-lived mutants simultaneously, under the same conditions. As demonstrated previously, *ife-2* (**A**), *clk-1* (**B**), *sod-2* (**C**), *eat-2* (**D**), *osm-5* (**E**), *nuo-6* (**F**), *isp-1* (**G**), *daf-2* (**H**) and *glp-1* (**I**) mutants all show increased lifespan compared to wild-type control worms. The length of lifespan extension varied from 4 days (25% increase) in *ife-2* worms to 22 days (138% increase) in *daf-2* worms (**J**). For *glp-1* worms and their wild-type controls, worms were grown at 25°C during development and were shifted to 20 °C at adulthood. Three biological replicates per strain were performed. Significance was determined using the log-rank test.

**Figure S2.**
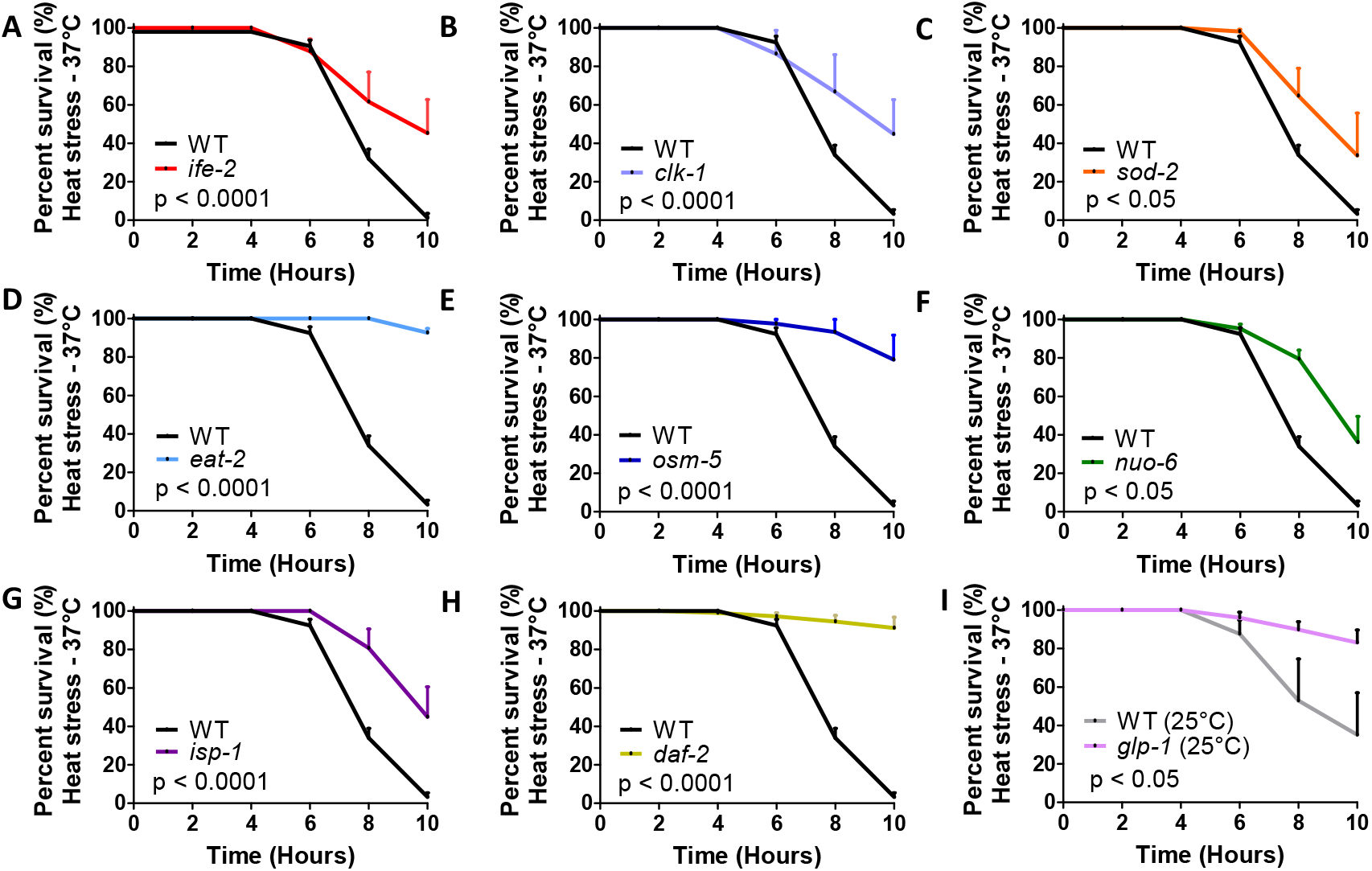
All long-lived mutants have increased resistance to heat stress. *ife-2* (**A**), *clk-1* (**B**), *sod-2* (**C**), *eat-2* (**D**), *osm-5* (**E**), *nuo-6* (**F**), *isp-1* (**G**), *daf-2* (**H**) and *glp-1* (**I**) worms show increased resistance to 37°C heat stress compared to wild-type worms. For *glp-1* worms and their wild-type controls, worms were grown at 25°C during development and were shifted to 20 °C at adulthood. Error bars represent SEM. Three biological replicates per strain were performed. Significance was determined using a repeated measures ANOVA.

**Figure S3.**
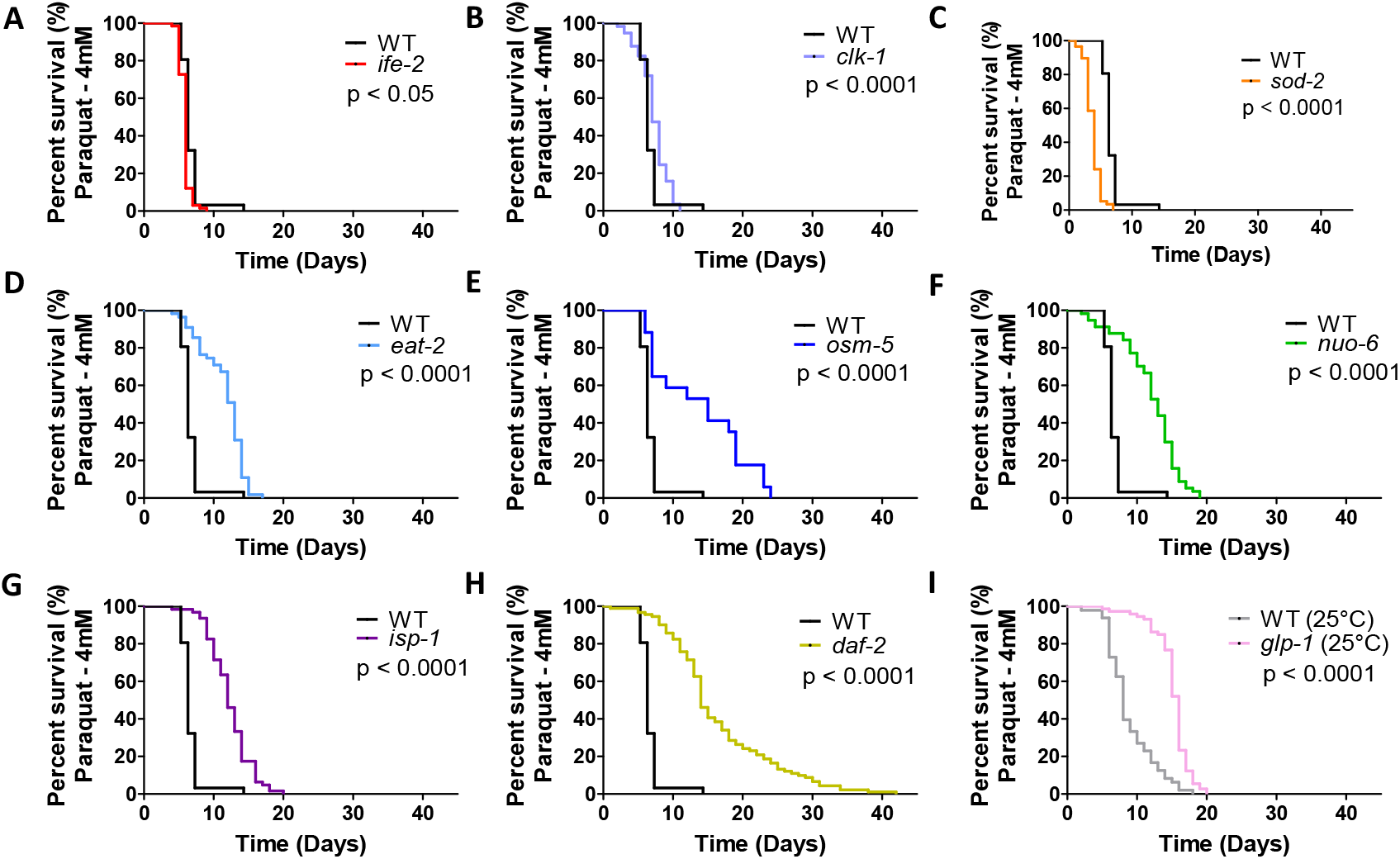
Most but not all long-lived mutants exhibit increased resistance to chronic oxidative stress resulting from exposure to paraquat. Resistance to chronic oxidative stress was measured by monitoring survival on plates containing 4 mM paraquat. *ife-2* (**A**) and *sod-2* (**C**) mutants show less resistance to paraquat compared to wild-type worms. *clk-1* (**B**), *eat-2* (**D**), *osm-5* (**E**), *nuo-6* (**F**), *isp-1* (**G**), *daf-2* (**H**), and *glp-1* (**I**) worms show increased resistance to paraquat compared to wild-type worms. All six of the longest lived strains show increased resistance to paraquat. For *glp-1* mutants and their wild-type controls, worms were grown at 25°C during development and were shifted to 20 °C at adulthood. Three biological replicates per strain were performed. Significance was determined using the log-rank test.

**Figure S4.**
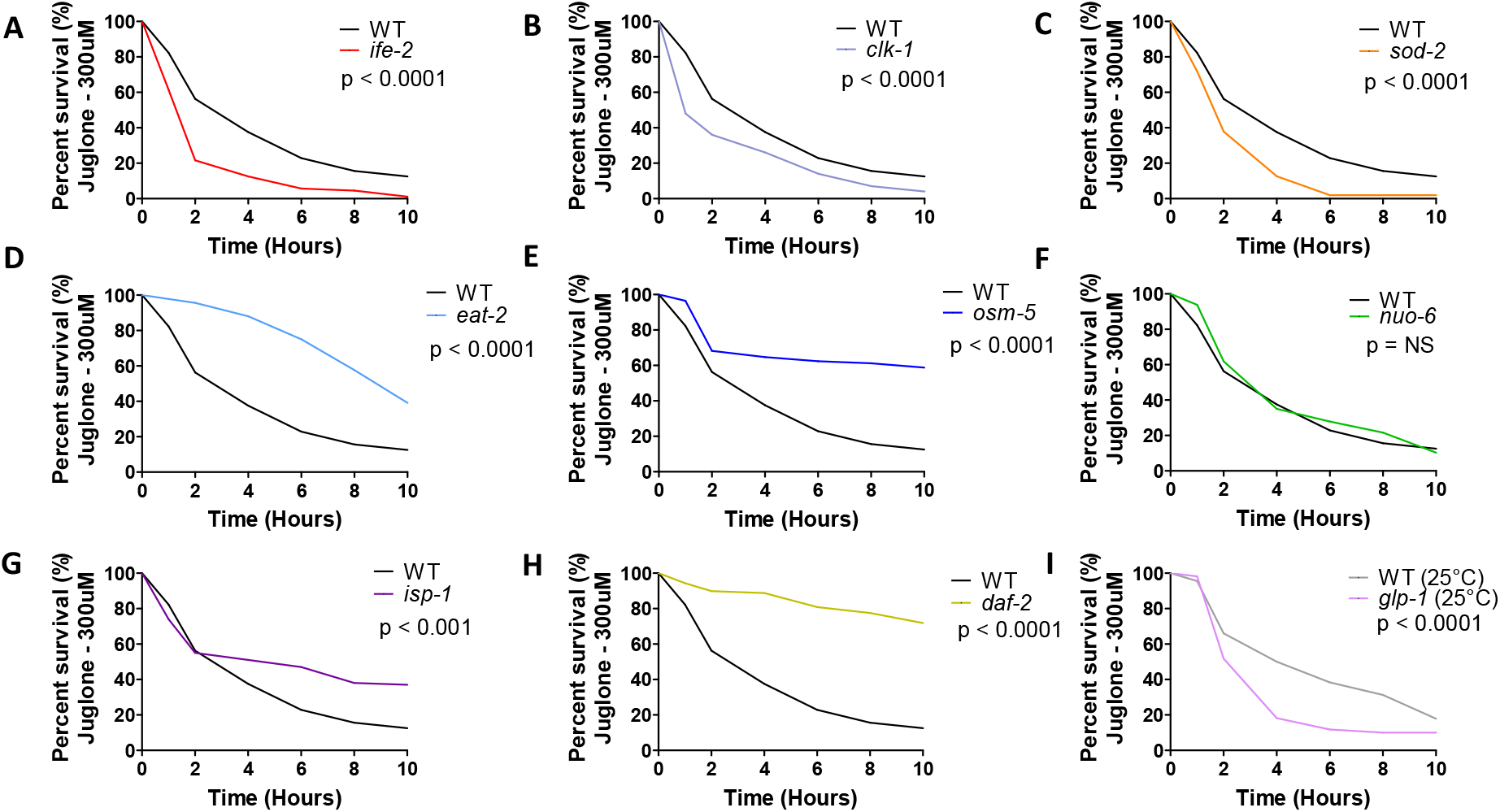
Long-lived mutants show variable resistance to acute oxidative stress resulting from exposure to juglone. Resistance to acute oxidative stress was measured by monitoring survival on plates containing 300 μM juglone. *ife-2* (**A**), *clk-1* (**B**), *sod-2* (**C**) and *glp-1* (**I**) mutants show less resistance to juglone compared to wild-type worms. *eat-2* (**D**), *osm-5* (**E**), *isp-1* (**G**), and *daf-2* (**H**) worms show increased resistance to juglone compared to wild-type worms, while *nuo-6* (**F**) worms showed no difference. For *glp-1* mutants and their wild-type controls, worms were grown at 25°C during development and were shifted to 20 °C at adulthood. Three biological replicates per strain were performed. Significance was determined using the log rank test.

**Figure S5.**
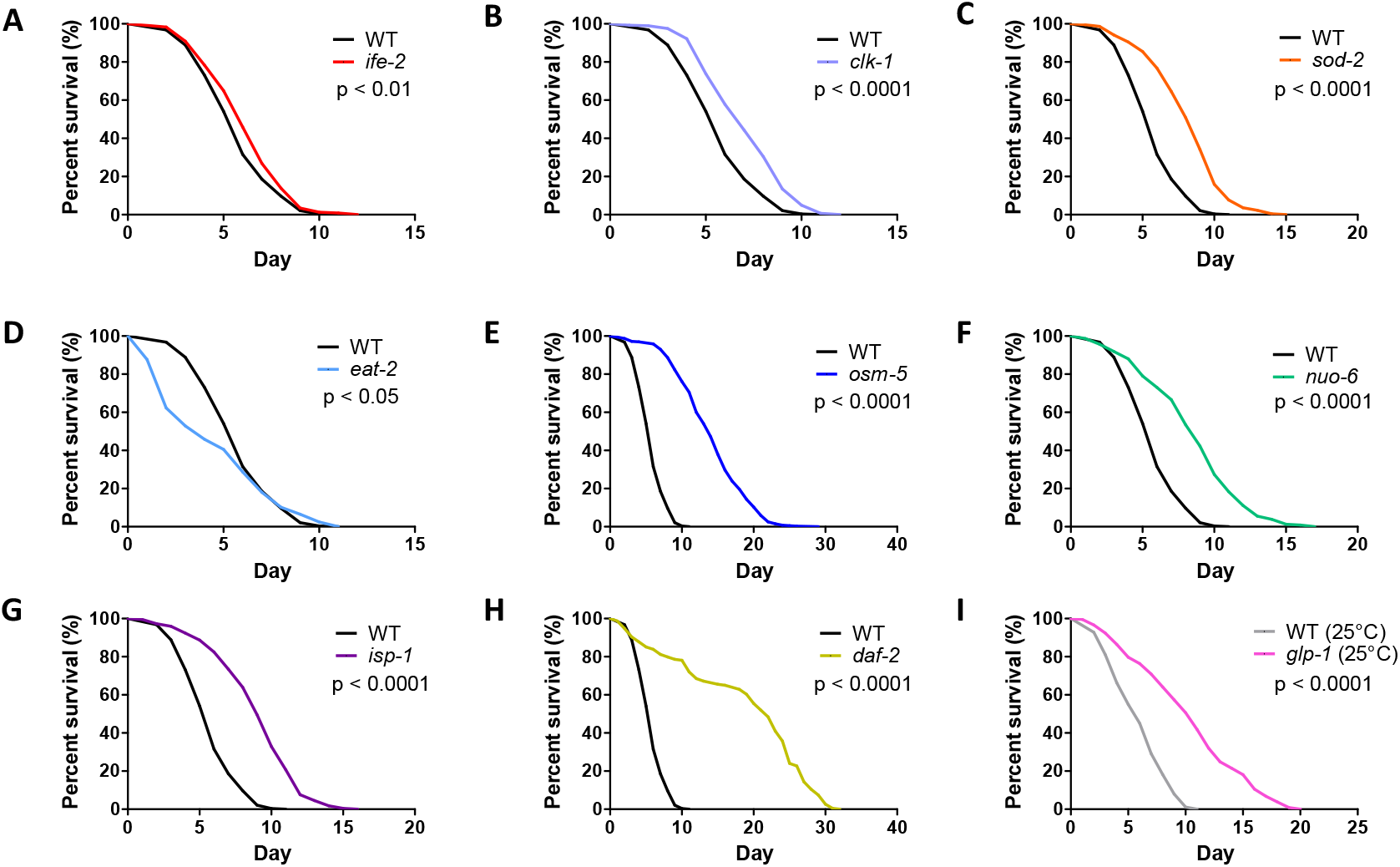
Most but not all long-lived mutants exhibit increased resistance to bacterial pathogen stress. Resistance to bacterial pathogen stress was assessed by exposing worms to *Pseudomonas aeruginosa* strain PA14 in a slow kill assay. While *eat-2* (**D**) worms show less resistance to PA14 compared to wild-type worms, *ife-2* (**A**), *clk-1* (**B**), *sod-2* (**C**), *osm-5* (**E**), *nuo-6* (**F**), *isp-1* (**G**), *daf-2* (**H**) and *glp-1* (**I**) mutants all show increased resistance to PA14 compared to wild-type worms. For *glp-1* worms and their wild-type controls, worms were grown at 25°C during development and were shifted to 20 °C at adulthood. Three biological replicates per strain were performed. Significance was determined using the log-rank test.

**Figure S6.**
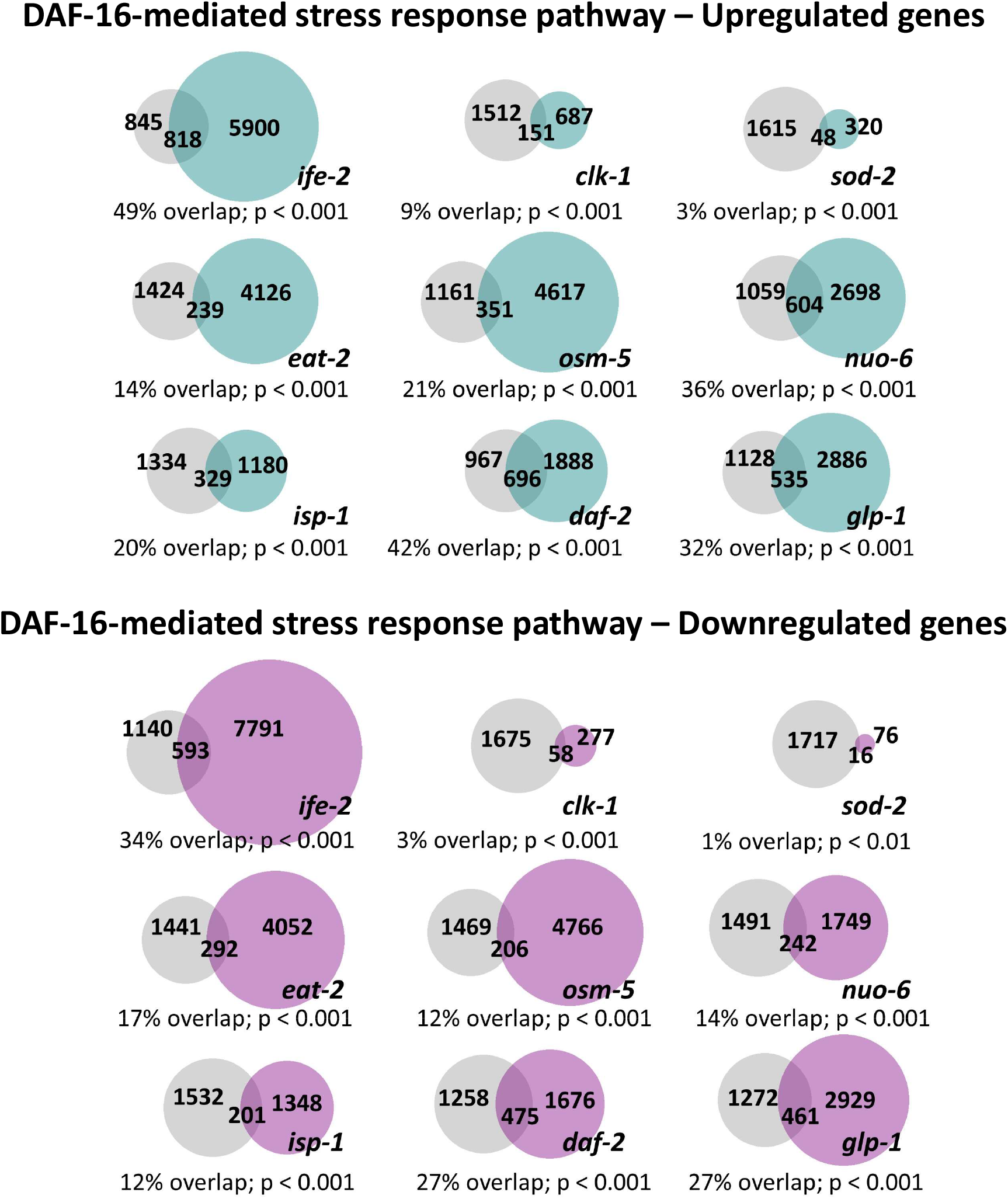
Long-lived genetic mutants exhibit significant modulation of genetic targets of DAF-16-mediated stress response pathway. Upregulated genetic targets of DAF-16 (grey) overlap with upregulated genes in long-lived mutants (teal), ranging from 3% to 49% overlap (**Top**). Downregulated genetic targets of DAF-16 (grey) overlap with downregulated genes in long-lived mutants (lavender), ranging from 1% to 34% overlap (**Bottom**).

**Figure S7.**
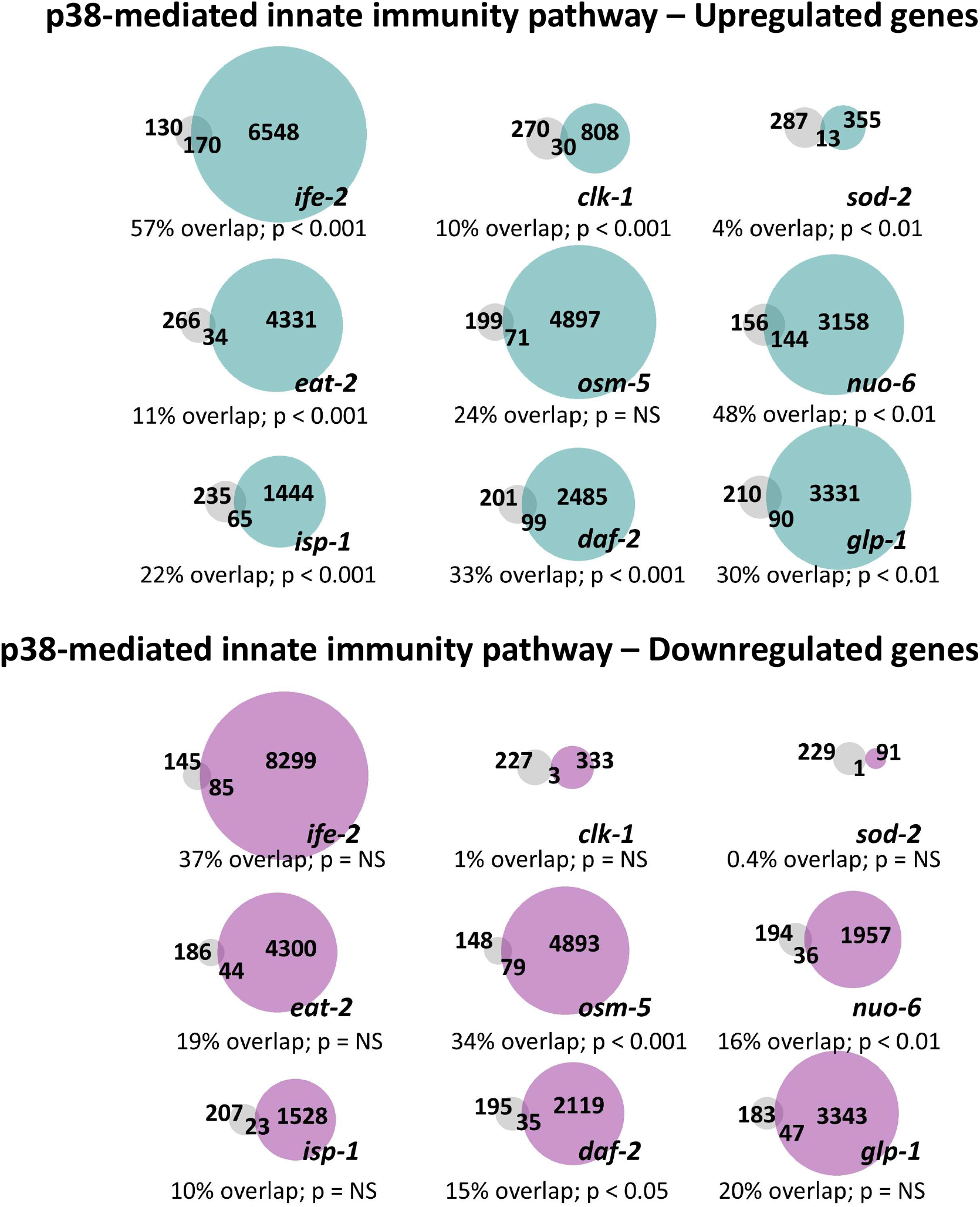
Long-lived genetic mutants exhibit significant modulation of genetic targets of the p38-mediated innate immunity pathway. Upregulated genetic targets of p38-mediated innate immunity pathway (grey) overlap with upregulated genes in long-lived mutants (teal), ranging from 4% to 57% overlap (**Top**). Downregulated genetic targets of p38-mediated pathway (grey) overlap with downregulated genes in long-lived mutants (lavender), ranging from 0.4% to 37% overlap (**Bottom**).

**Figure S8.**
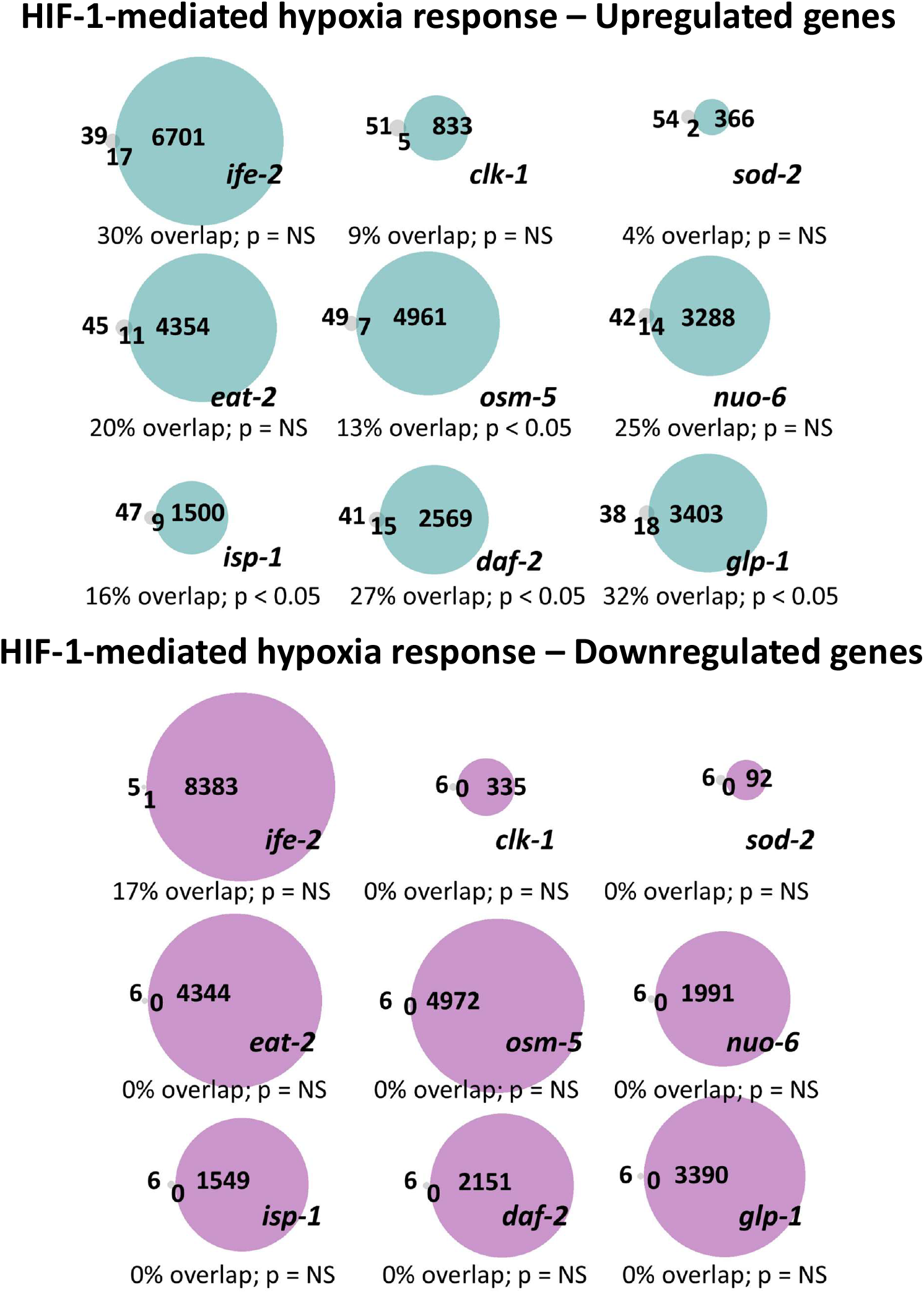
Long-lived genetic mutants exhibit significant modulation of genetic targets of HIF-1-mediated hypoxia pathway. Upregulated genetic targets of HIF-1 (grey) overlap with upregulated genes in long-lived mutants (teal), ranging from 4% to 32% overlap (**Top**). Downregulated genetic targets of HIF-1 (grey) overlap with downregulated genes in long-lived mutants (lavender), ranging from 0% to 17% overlap (**Bottom**).

**Figure S9.**
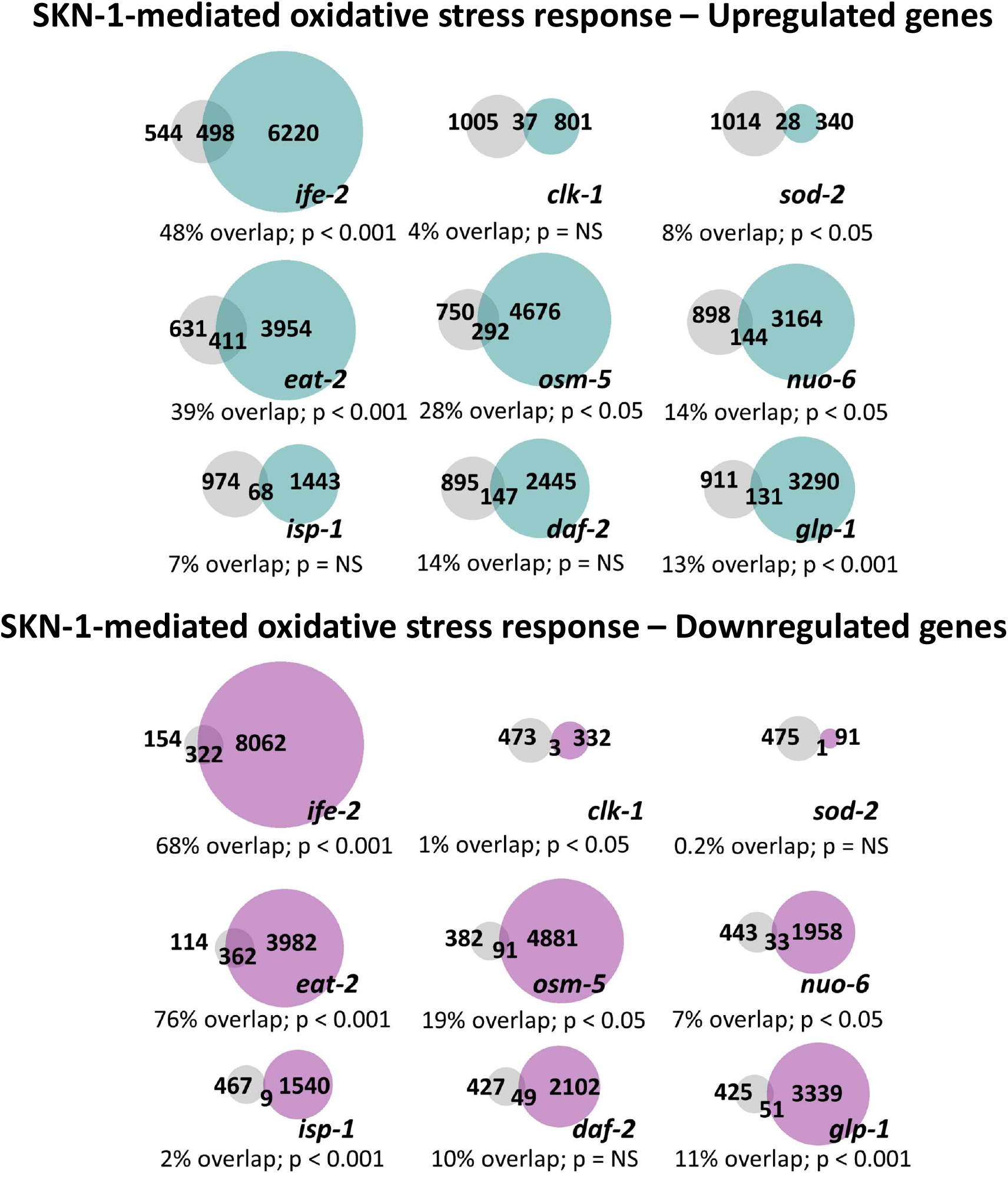
Long-lived genetic mutants exhibit significant modulation of genetic targets of SKN-1-mediated oxidative stress response pathway. Upregulated genetic targets of SKN-1 (grey) overlap with upregulated genes in long-lived mutants (teal), ranging from 4% to 48% overlap (**Top**). Downregulated genetic targets of SKN-1 (grey) overlap with downregulated genes in long-lived mutants (lavender), ranging from 0.2% to 76% overlap (**Bottom**).

**Figure S10.**
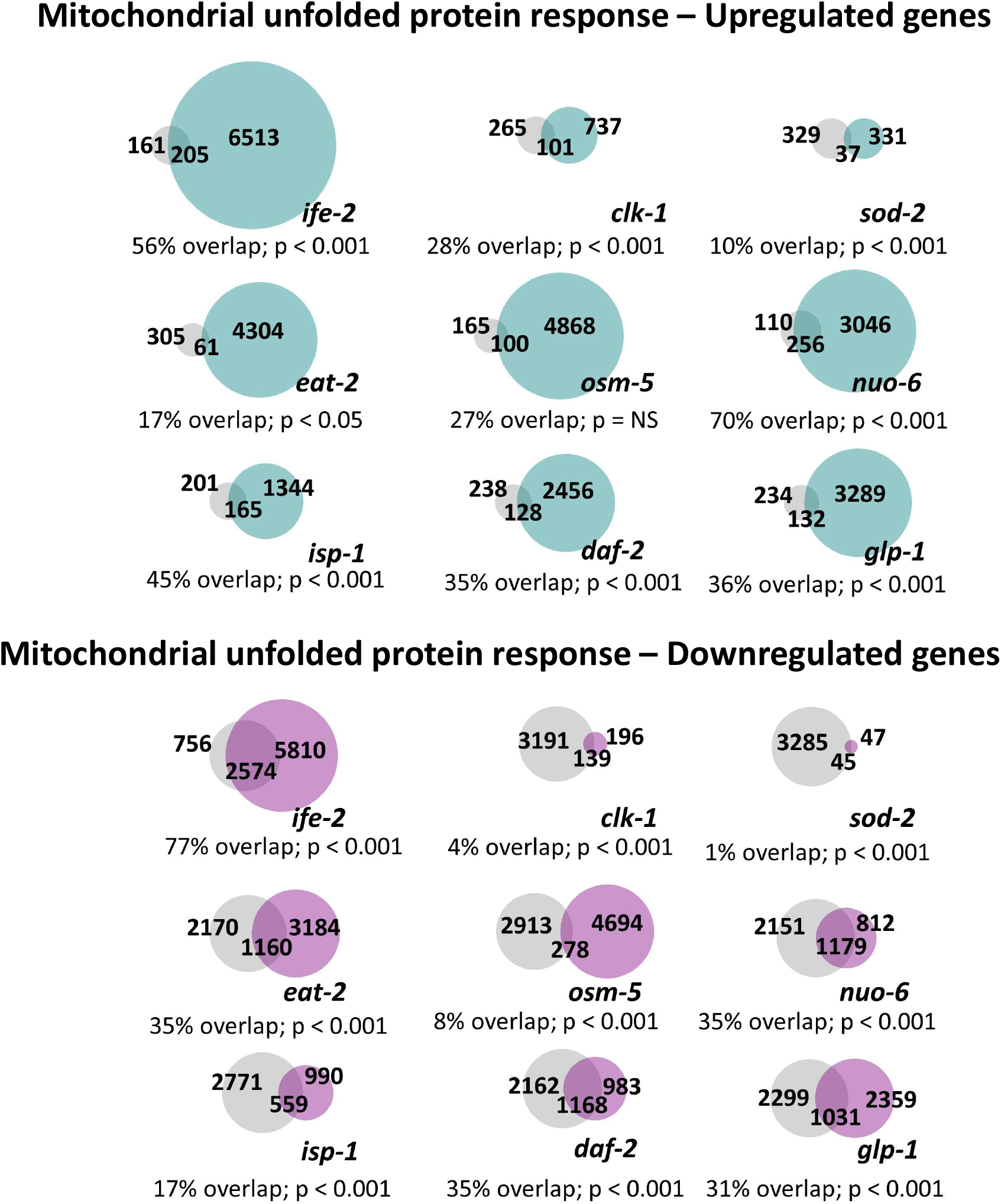
Long-lived genetic mutants exhibit significant modulation of genetic targets of the mitochondrial unfolded protein response pathway. Upregulated genetic targets of the mitoUPR pathway (grey) overlap with upregulated genes in long-lived mutants (teal), ranging from 10% to 70% overlap (**Top**). Downregulated genetic targets of mitoUPR pathway (grey) overlap with downregulated genes in long-lived mutants (lavender), ranging from 1% to 77% overlap (**Bottom**).

**Figure S11.**
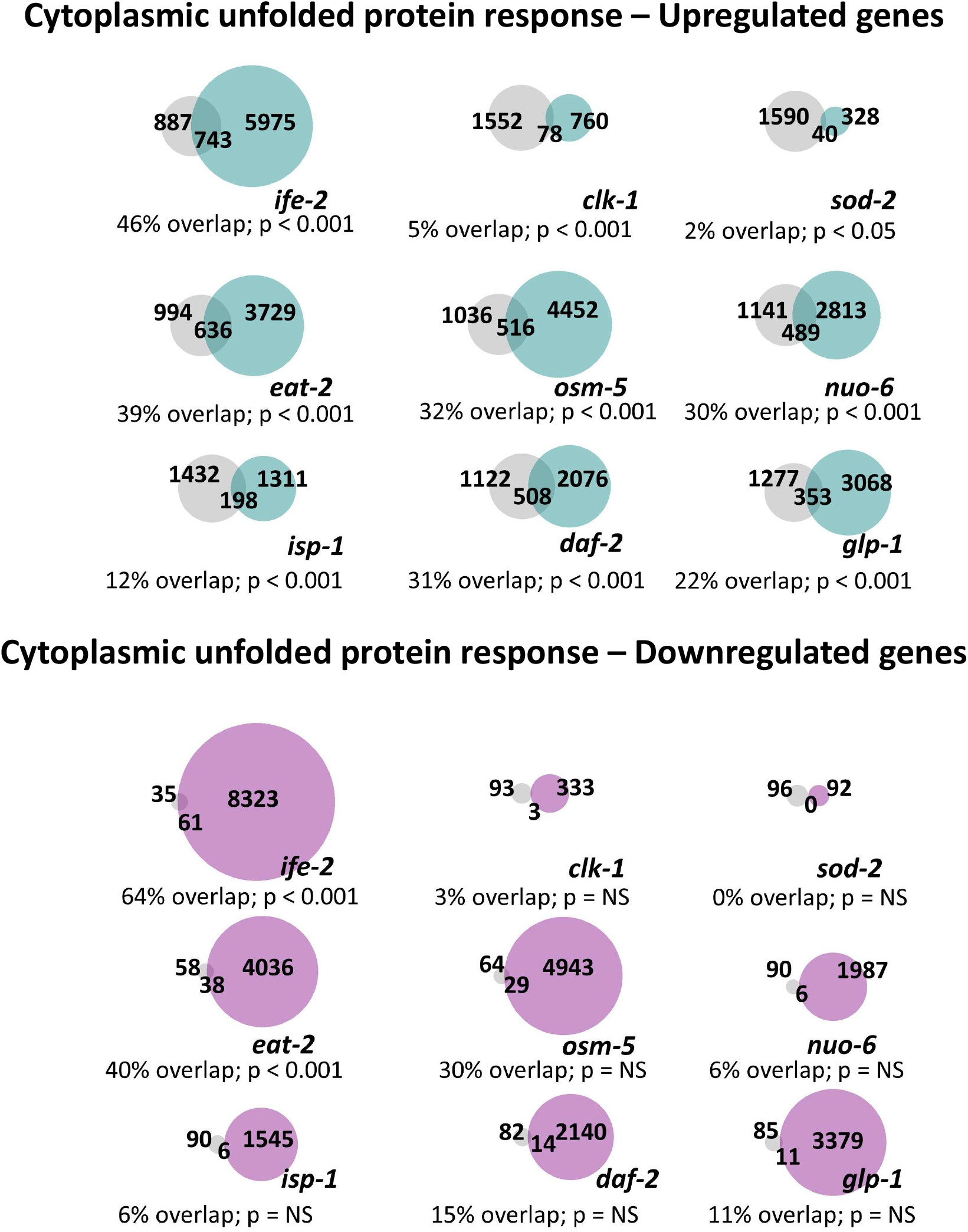
Long-lived genetic mutants exhibit significant modulation of genetic targets of cytoplasmic unfolded protein response pathway. Upregulated genetic targets of cyto-UPR pathway(grey) overlap with upregulated genes in long-lived mutants (teal), ranging from 2% to 46% overlap (**Top**). Downregulated genetic targets of the cyto-UPR pathway (grey) overlap with downregulated genes in long-lived mutants (lavender), ranging from 0% to 64% overlap (**Bottom**).

**Figure S12.**
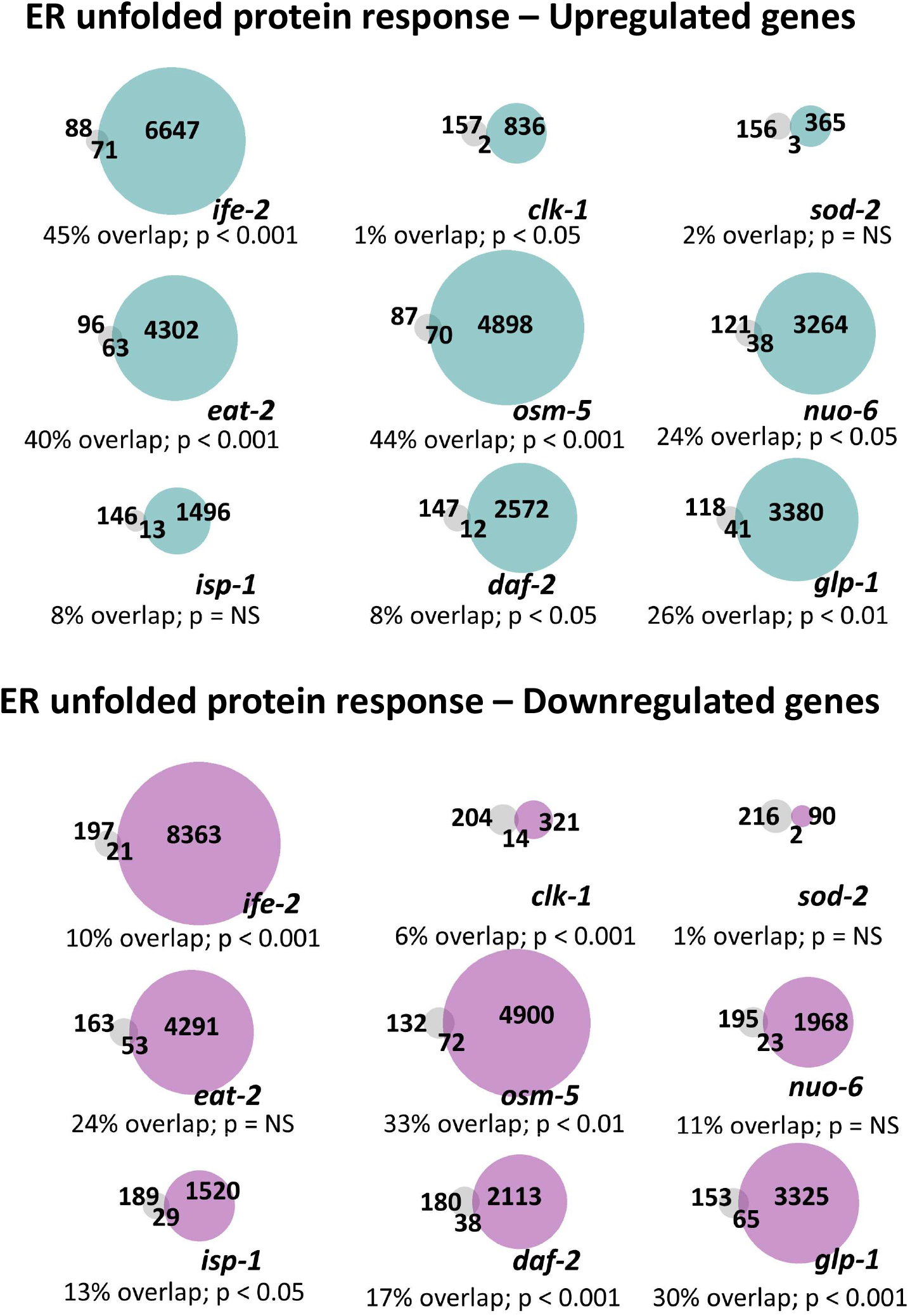
Long-lived genetic mutants exhibit significant modulation of genetic targets of the ER unfolded protein response pathway. Upregulated genetic targets of ER-UPR pathway (grey) overlap with upregulated genes in long-lived mutants (teal), ranging from 1% to 45% overlap (**Top**). Downregulated genetic targets of ER-UPR pathway (grey) overlap with downregulated genes in long-lived mutants (lavender), ranging from 1% to 33% overlap (**Bottom**).

**Figure S13.**
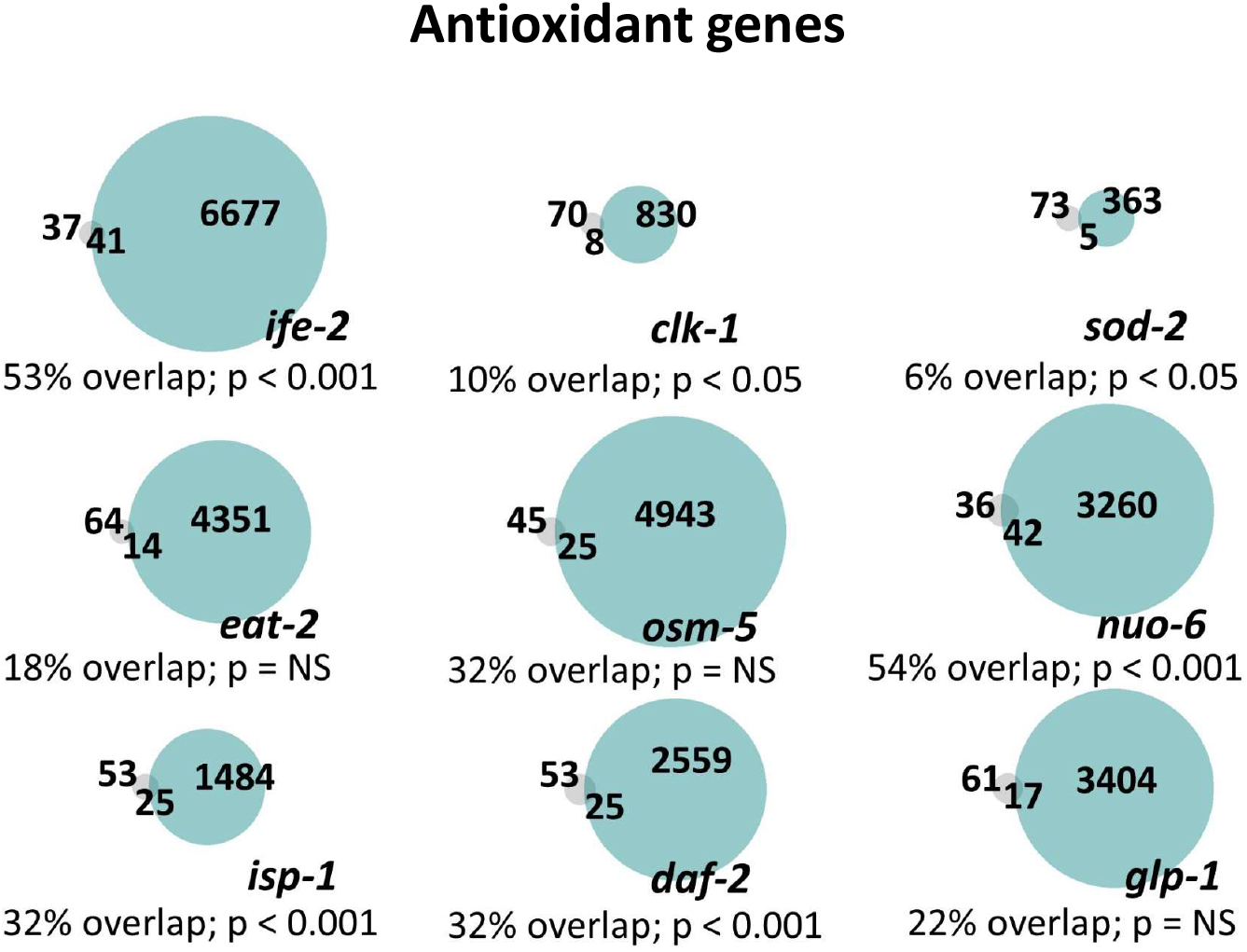
Long-lived genetic mutants exhibit significant modulation of antioxidant genes. Antioxidant genes (grey) overlap with upregulated genes in long-lived mutants (teal), ranging from 6% to 54% overlap.

**Figure S14.**
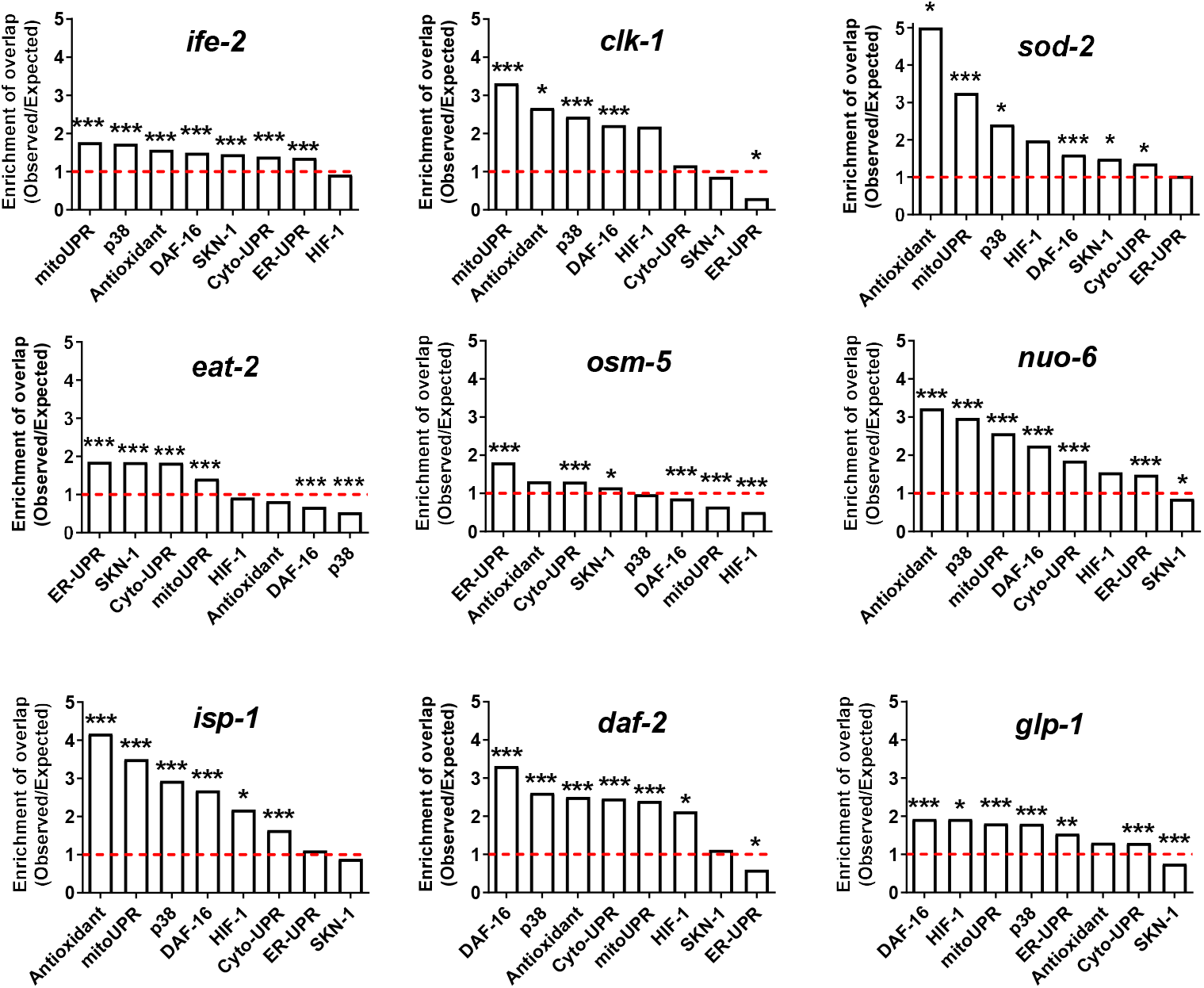
Relative contribution of different stress response pathways in long-lived mutants. Gene expression in the long-lived mutant strains was examined by RNA sequencing (RNA-seq) of six biological replicate per genotype of pre-fertile day 1 young adult worms. Differentially expressed genes that are significantly upregulated in the long-lived mutant strains were compared to genes that are upregulated by activation of different stress response pathways including the DAF-16-mediated stress response (DAF-16), the p38-regulated innate immune signaling pathway (p38), the HIF-1-mediated hypoxia response (HIF-1), the SKN-1-mediated oxidative stress response (SKN-1), the mitochondrial unfolded protein response (mitoUPR), the cytoplasmic unfolded protein response (Cyto-UPR), the ER unfolded protein response (ER-UPR), and antioxidant gene expression (Antioxidant). For each stress pathway, the ratio of the observed number of overlapping genes with the long-lived mutant to the expected number of overlapping genes if picked randomly was determined. All of the long-lived mutants showed a significant enrichment of genes involved in at least three stress response pathways. For each mutant, the stress response pathways are arranged in descending order of observed overlapping genes/expected overlapping genes.

**Figure S15.**
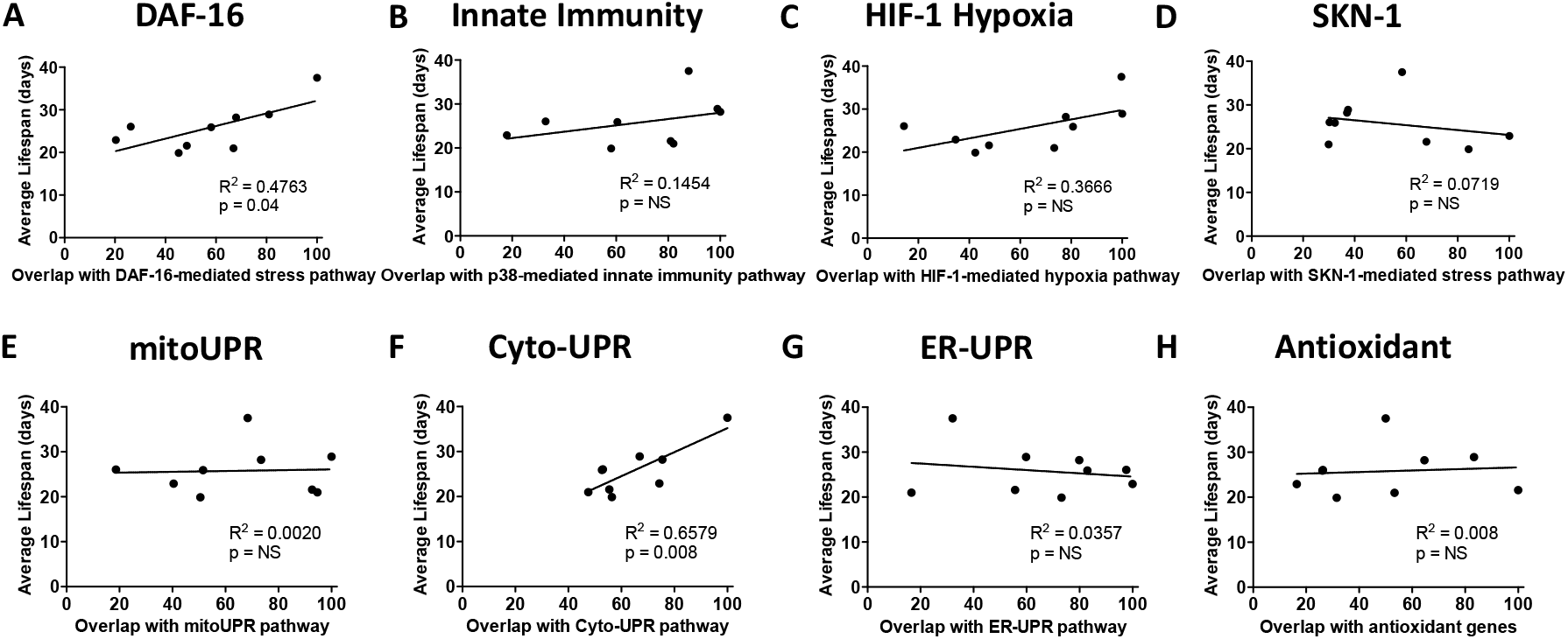
Degree of overlap with genes in the cytoplasmic unfolded protein response pathway and DAF-16-mediated stress response pathway are correlated with lifespan. While there is a significant degree of overlap between genes involved in stress response pathways and genes upregulated in long-lived mutants, there is only a significant correlation between the degree of overlap and the magnitude of lifespan extension for the DAF-16-mediated stress response pathway and the cytoplasmic unfolded protein response.

**Figure S16.**
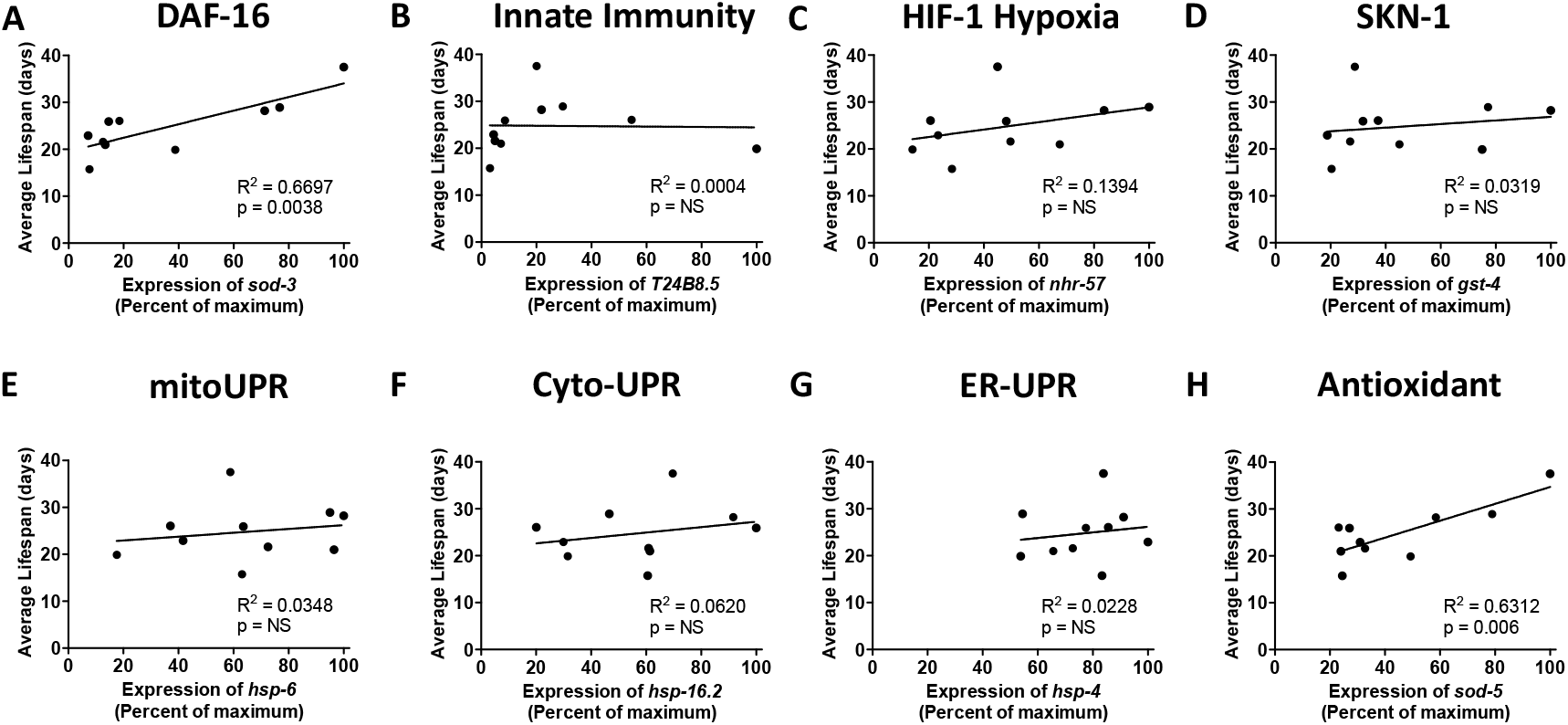
Expression levels of inducible superoxide dismutase genes are correlated with magnitude of lifespan extension. Comparing the magnitude of gene upregulation of target genes of different stress response pathways to the magnitude of lifespan extension in nine long-lived mutants revealed that the expression levels of target genes from the DAF-16-mediated stress response pathway (*sod-3*) and antioxidant defense pathway (*sod-5*) are significantly correlated with the magnitude of lifespan extension.

**Figure S17.**
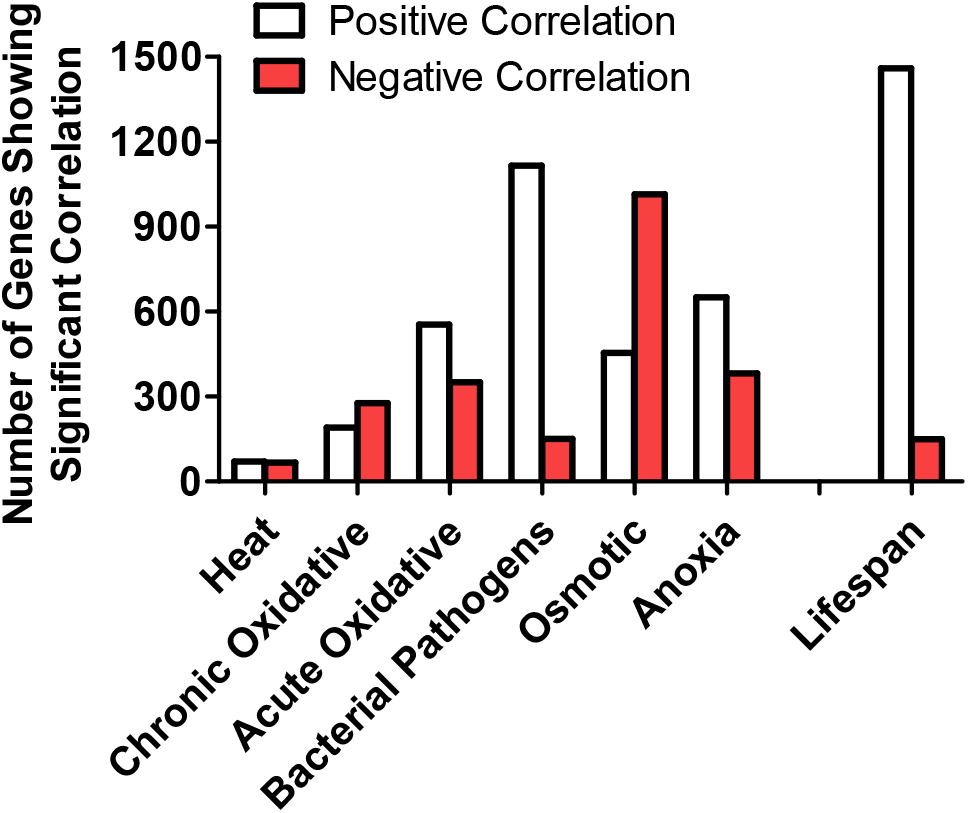
Genes correlated with resistance to different external stressors. The expression levels of all of the genes in the genome were determined by RNA sequencing in nine long-lived mutants. These expression levels were compared to the survival of these strains when exposed to different types of stress to determine which genes are correlated with stress resistance. There are numerous genes that exhibit either a positive or negative correlation with each type of stress resistance indicating a strong influence of genetics on stress resistance. Complete lists of genes that are significantly correlated with each type of stress resistance can be found in **Table S3**.

**Figure S18.**
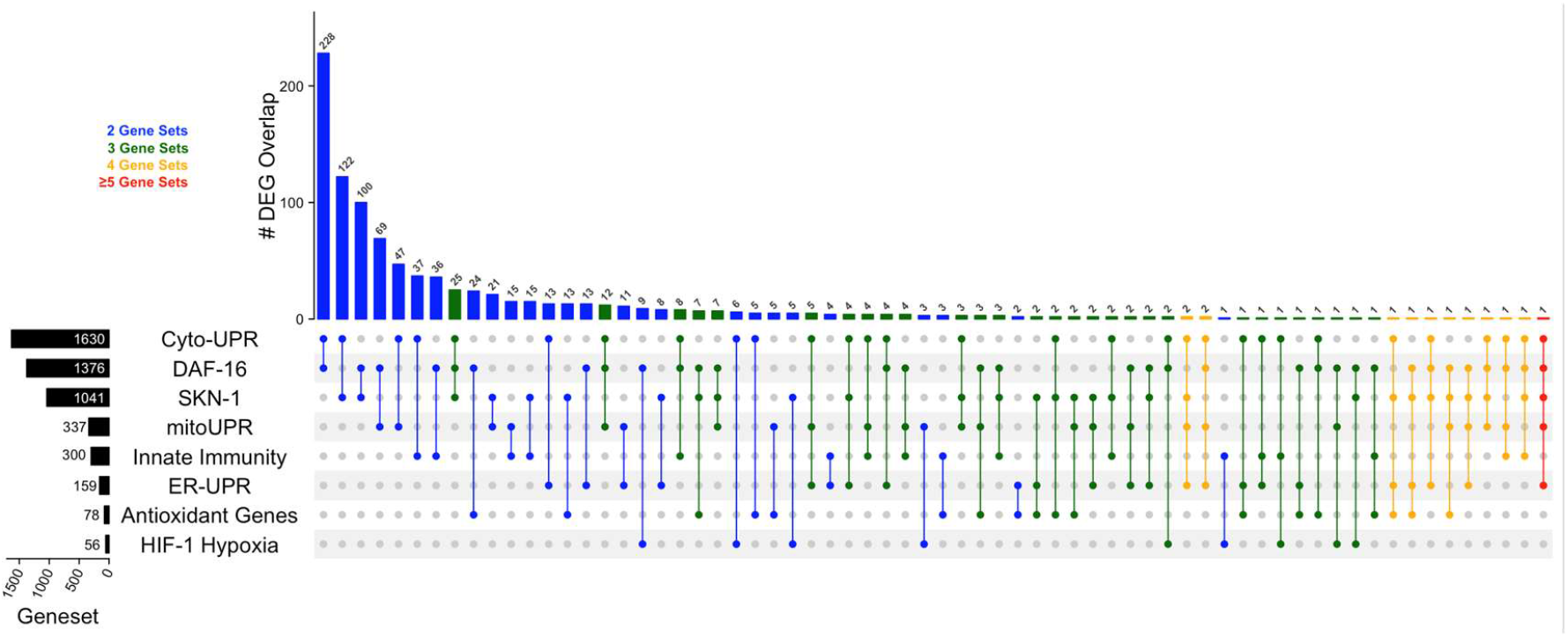
Genes upregulated by activation of stress response pathways exhibit partial overlap with other stress response pathways. This inclusive UpSetR plot compares genes that are upregulated by activation of each stress response pathway. Stress response pathways are listed on the left in order of number of upregulated genes from top to bottom. The number of genes that are upregulated by activation of each pathway is indicated by black bars and associated numbers. On the main plot, the height of each bar and the number above the bar indicate the number of genes in common between the gene sets indicated by the dots below the plot. The complete lists of genes that are upregulated by activation of each stress response pathway can be found in **Table S2**. Cyto-UPR = cytoplasmic unfolded protein response; DAF-16 = DAF-16-mediated stress response pathway; SKN-1= SKN-1-mediated oxidative stress response pathway; mitoUPR = mitochondrial unfolded protein response; Innate immunity = p38-mediated innate immune pathway; ER-UPR = endoplasmic reticulum unfolded protein response; HIF-1 Hypoxia = HIF-1-mediated hypoxia pathway.

**Figure S19.**
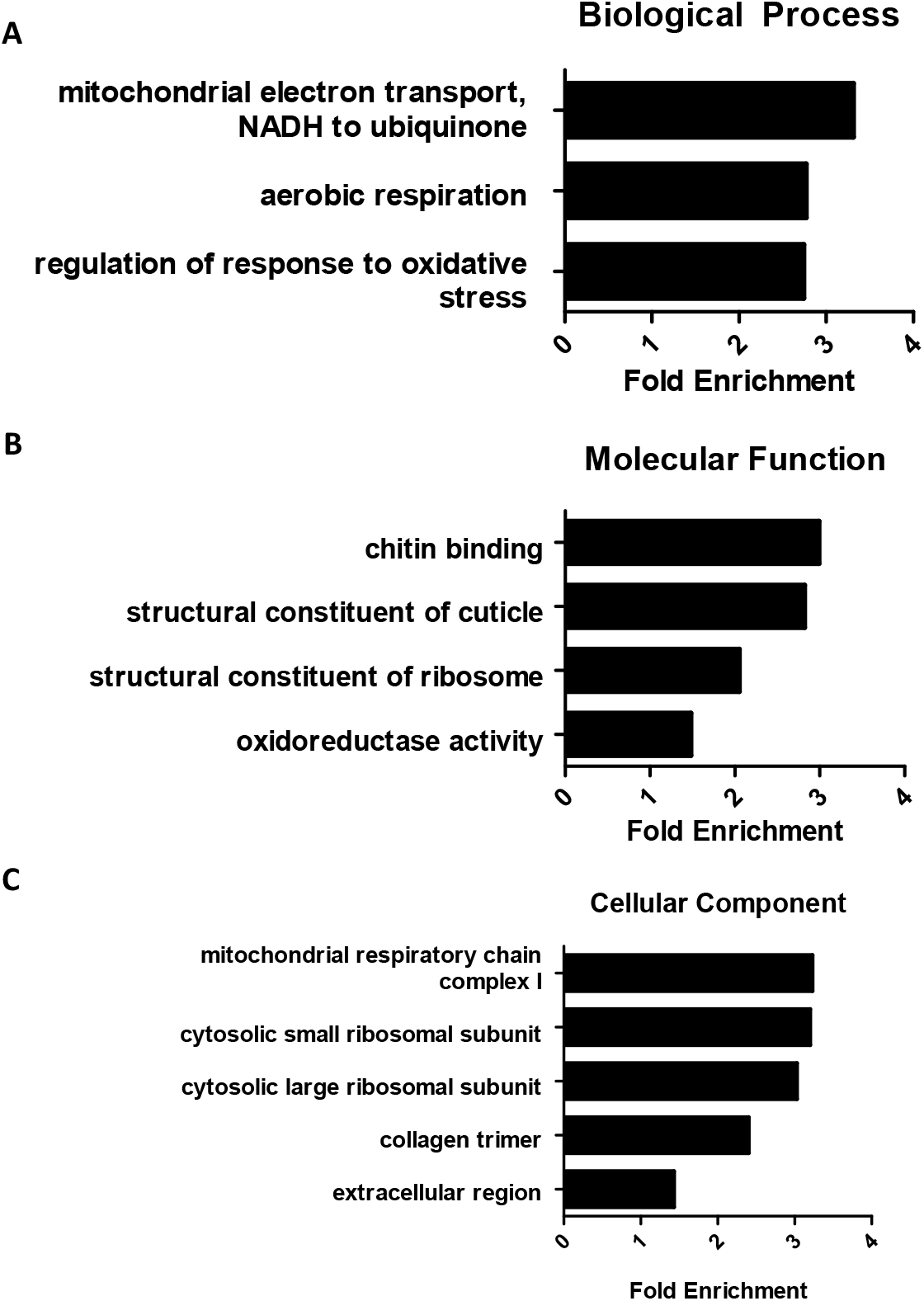
Enrichment analysis of genes correlated with resistance to stress reveals a role for mitochondrial function and structures providing a barrier to the environment in stress resistance. To determine the functional classes that are overrepresented in genes correlated with at least one type of stress, we used the statistical overrepresentation test of Gene Ontology (GO) terms with PANTHER Classification System (version 16.0) and compared the genes to the *C. elegans* genome. PANTHER recognized 2997 genes and found 3 significantly enriched classes for Biological Process (**A**), 4 significantly enriched classes for Molecular Function (**B**), and 5 significantly enriched classes for Cellular Component (**C**).

**Figure S20.**
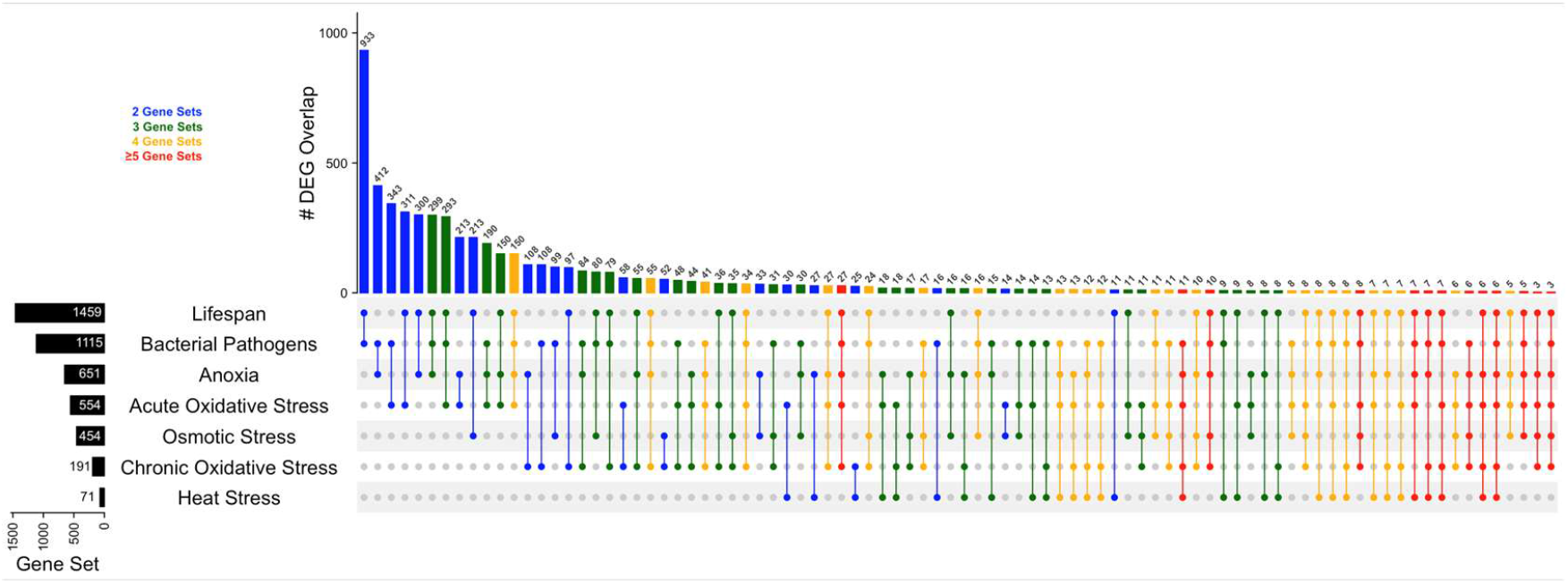
Highly significant overlap between genes the are correlated with resistance to stress and genes that are correlated with lifespan extension. Inclusive UpSetR plot comparing genes correlated with lifespan and genes correlated with different types of stress resistance including bacterial pathogens (*P. aeruginosa*), anoxic stress (72-96 hours), acute oxidative stress (300 μM juglone), chronic oxidative stress (4 mM paraquat), osmotic stress (450-500 mM NaCl), and heat stress (37°C). Black bars on the left indicate the number of genes in each gene set. Full lists of genes correlated with each type of stress resistance can be found in **Table S3**. Bars indicate the number of genes in common between the gene sets indicated by the dots below the plot. The exact number of genes in each intersection is shown above the graph. Intersections involving two gene sets are shown in blue. Intersections involving 3, 4 or 5-6 gene sets are shown in green, yellow or red, respectively.

**Figure S21.**
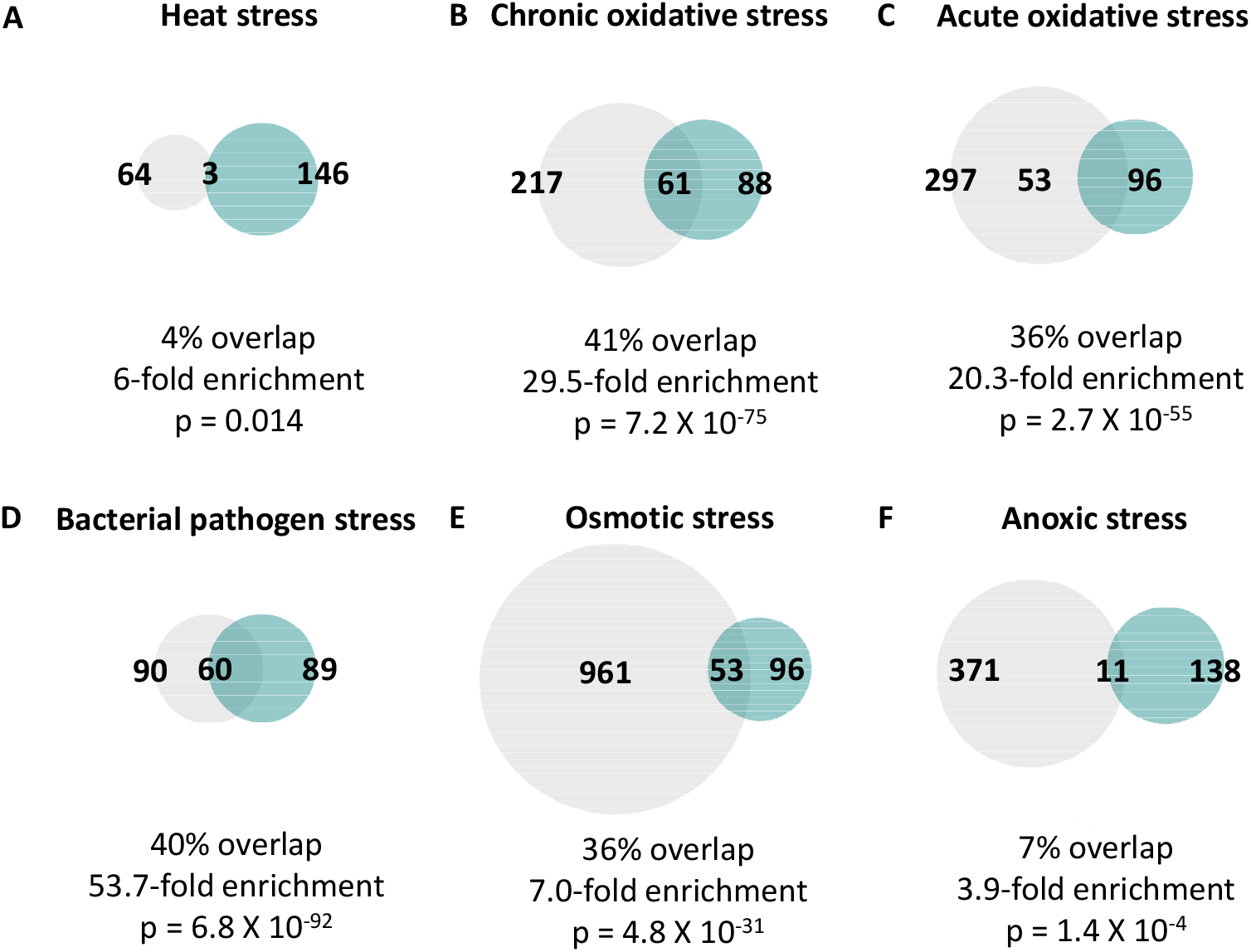
Highly significant overlap between genes the are negatively correlated with resistance to stress and genes that are negatively correlated with lifespan extension. In examining the degree of overlap between genes that have a significant negative correlation with lifespan and genes that exhibit a significant negative correlation with resistance to stress, we observed a significant degree of overlap with each of the six types of stress resistance that we examined including heat stress (**A**; 37°C), chronic oxidative stress (**B**; 4 mM paraquat), acute oxidative stress (**C**; 300 μM juglone), bacterial pathogen stress (**D**; *P. aeruginosa*), osmotic stress (**E**; 450-500 mM NaCl), and anoxia stress (**F**; 72-96 hours). The degree of enrichment ranged from 3.9 fold up to 53.7 fold with percent overlap between 4% and 41%. The most highly significant overlap with genes negatively correlated with lifespan was with genes negatively correlated with bacterial pathogen stress survival.

**Figure S22.**
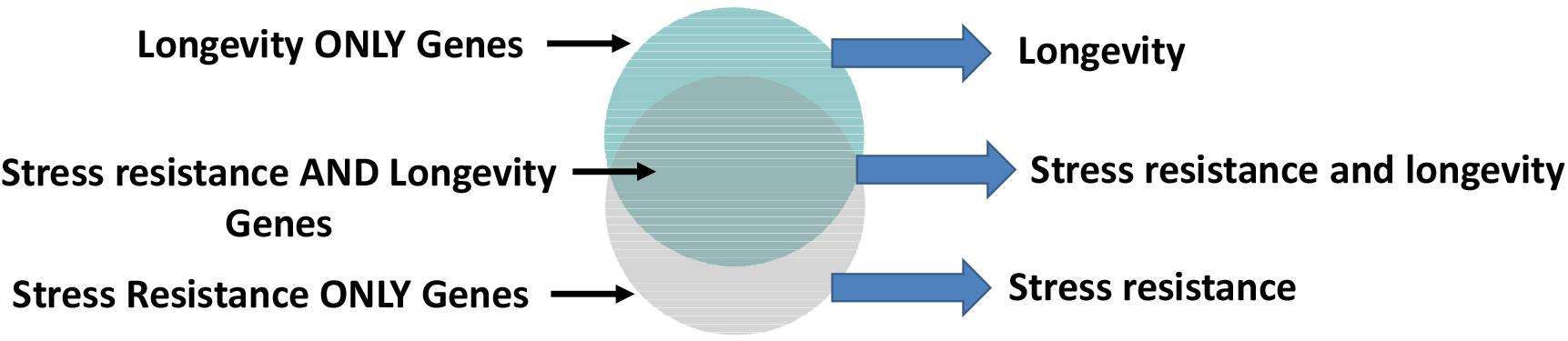
Model for relationship between stress resistance and lifespan. The genetic factors contributing to stress resistance and lifespan share a high degree of overlap, but there are also groups of genes that contribute only to one phenotype or the other. Modulating genes that contribute to both stress resistance and longevity will affect both phenotypes and account for the significant correlation that is observed between these two phenotypes. Modulating “Longevity ONLY genes” or “Stress resistance ONLY genes” affects longevity or stress resistance independently of the other, thereby allowing for these phenotypes to be experimentally dissociated.

**Table S1.** Summary of stress resistance in long-lived mutant strains. “=“ indicates no significant difference from wild-type “-” indicates decreased compared to wild-type; “+” indicates increased compared to wild-type; “++” indicates markedly increased compared to wild-type.

| Strain | Lifespan Extension | Acute oxidative Stress | Chronic oxidative Stress | Heat Stress | Anoxic Stress | Osmotic Stress | PA14 Stress |
| --- | --- | --- | --- | --- | --- | --- | --- |
| *ife-2* | 26.3% | - | - | + | = | - | = |
| *clk-1* | 33.4% | - | + | + | = | = | + |
| *sod-2* | 37.2% | - | - | + | = | = | + |
| *eat-2* | 45.6% | + | + | ++ | = | -- | - |
| *osm-5* | 65.4% | ++ | ++ | ++ | ++ | ++ | ++ |
| *nuo-6* | 79.2% | = | + | + | - | ++ | + |
| *isp-1* | 83.8% | + | + | + | = | ++ | + |
| *glp-1* | 89.2% | - | + | + | + | ++ | + |
| *daf-2* | 138.4% | ++ | ++ | ++ | ++ | ++ | ++ |

## Notes

### Competing Interest Statement

The authors have declared no competing interest.

